# EMILIN1 emerges as a TGFβ/SETDB1-regulated secreted biomarker in Duchenne Muscular Dystrophy

**DOI:** 10.1101/2025.10.11.681604

**Authors:** Maeva Zamperoni, Laura Muraine, Alice Granados, Anne Bigot, Valentin Petit, Mona Bensalah, Jessica Ohana, Véronique Legros, Ekaterina Boyarchuk, Johanna Bruce, Guillaume Chevreux, Véronique Joliot, Elisa Negroni, Maryline Moulin, Capucine Trollet, Slimane Ait-Si-Ali

**Affiliations:** Université Paris Cité, CNRS, Epigenetics and Cell Fate, UMR7216, F-75013 Paris, France; Sorbonne Université, Inserm, Institut de Myologie, Centre de Recherche en Myologie, Paris, France; Université Paris Cité, CNRS, Institut Jacques Monod, UMR7592, F-75013 Paris, France; Proteom’IC facility, Université Paris Cité, CNRS, INSERM, Institut Cochin, F-75014 Paris, France

## Abstract

Duchenne Muscular Dystrophy (DMD) is an incurable muscle-wasting disorder characterized by chronic membrane damage, inflammation, and progressive fibrosis. Fibrosis in DMD is driven by sustained TGFβ signaling, which promotes extracellular matrix (ECM) accumulation. We previously showed that SETDB1 sustains the TGFβ-induced fibrotic response in DMD myotubes. Here, we further show that SETDB1 modulates the TGFβ-induced secretome, particularly by regulating ECM-related proteins. Comparison of the basal secretome from DMD patient-derived myotubes and healthy controls revealed a distinct disease-specific profile. Integrating both secretome analyses, we identified EMILIN1, an ECM glycoprotein not previously studied in skeletal muscle, as a robust shared candidate; EMILIN1 is enriched in the DMD secretome, further upregulated by TGFβ, and downregulated upon SETDB1 depletion. We confirmed EMILIN1 overexpression in DMD patient muscle biopsies, validating its pathological relevance. Functionally, EMILIN1 depletion modulated myogenic differentiation and reduced expression of the fibrotic marker SERPINE1. These findings establish EMILIN1 as a novel secreted regulator of myogenesis and fibrosis, and implicate SETDB1 in shaping the TGFβ-dependent secretome in DMD. Our integrative proteomic approach provides new insights into the molecular drivers of impaired regeneration in DMD and highlights potential therapeutic targets.

## INTRODUCTION

Skeletal muscle is a highly organized structure composed of various tissues, including blood and lymphatic vessels, contractile muscle fibers, and connective tissue (Dave HD, 2025). The cooperation between these tissues guarantees the regenerative ability of skeletal muscle upon injury. Normally, after mechanical trauma, exposure to toxins or infections, skeletal muscle retains the ability to completely repair, thanks to a sequential and well-orchestrated series of events, referred as muscle regeneration. However, in conditions of disease, like in DMD, the inflammatory response becomes chronic, leading to fibrosis, defined as an excessive accumulation of ECM components, and to fat accumulation, which ultimately replaces functional muscle tissue with non-functional one (Cappellari et al., 2020; Serrano et al., 2011).

DMD is an X-chromosome-linked recessive muscle wasting disorder with a prevalence of approximately 1 in 3500 live male births (Mercuri & Muntoni, 2013). DMD is due to deleterious mutations in the *DMD* gene leading to the lack of a functional Dystrophin protein, and thus to a significant fragility of the sarcolemma. This weakening of the sarcolemma contributes to repeated cycles of muscle fiber injury and repair, ultimately resulting in progressive muscle degeneration. Over time, this degeneration leads to extensive muscle wasting (Blake et al., 2002; Kharraz et al., 2014). One of the key pathological features accompanying this process is fibrosis, which is sustained by the TGFβ pathway, overactivated in DMD (Choi et al., 2016).

We recently identified the lysine methyltransferase and epigenetic regulator SETDB1 as a modulator of TGFβ signaling in DMD (Granados et al., 2024). SETDB1 catalyzes histone H3 lysine 9 tri-methylation (H3K9me3), an epigenetic modification primarily associated with heterochromatin formation and gene repression, thereby contributing to the regulation of the cells’ functional states (e.g., stemness, proliferation, differentiation). In proliferating myoblasts, SETDB1 remains predominantly nuclear, where it prevents the expression of late differentiation genes through H3K9me3. Upon Wnt signaling activation at the onset of terminal differentiation, SETDB1 is exported to the cytoplasm, facilitating myotube formation (Beyer et al., 2016). TGFβ induces nuclear accumulation of SETDB1 in healthy myotubes, while SETDB1 is constantly accumulated in DMD myotube nuclei with intrinsic overactivated TGFβ pathway (Granados et al., 2024). Moreover, our previous results showed that SETDB1 regulates the expression of TGFβ- dependent secreted factors impacting fibrosis and myogenic differentiation. We demonstrated that the conditioned medium (CM) of TGFβ-treated myotubes impairs myoblasts differentiation and increases the expression of pro-fibrotic genes (Granados et al., 2024).

Here, we investigated how SETDB1 modulates the secretome of DMD myotubes using proteomics, with a focus on TGFβ-dependent targets. In parallel, we compared the secretome from DMD *versus* control myotubes under basal conditions to identify deregulated secreted factors associated with the disease context. Together, these complementary approaches allowed us to dissect both basal and TGFβ-induced alterations in the DMD secretome. We found that SETDB1 modulates the TGFβ-induced secretome, particularly by regulating the expression of secreted and ECM-related proteins, such as EMILIN1 (Elastin Microfibril Interface (EMI) Located proteIN 1). EMILIN1 is a glycoprotein primarily expressed in the ECM, where it contributes to various cellular processes through its C-terminal gC1q domain, which mediates interactions with integrins, promoting cell adhesion and migration, and through its EMI domain, involved in TGFβ processing (Colombatti et al., 2011). Studies performed in different contexts, like blood vessels (Zacchigna et al., 2006), skin psoriasis (Pivetta et al., 2022) and cancer (Honda et al., 2024) revealed an inhibitory role of EMILIN1 on TGFβ signaling.

The role and function of EMILIN1 in skeletal muscle have not been described yet. Here, we show that EMILIN1 is more highly expressed in DMD muscle cells and patient biopsies compared to healthy controls and may play a key role in regulating myogenic differentiation. Loss of EMILIN1 in myoblasts and myotubes results in increased expression of early myogenic markers, while late markers are reduced. Additionally, EMILIN1 modulates the expression of the fibrotic marker SERPINE1, which is regulated by both TGFβ and SETDB1. Notably, EMILIN1 depletion leads to a decrease in SERPINE1 expression. Our findings suggest that EMILIN1 is a TGFβ- and SETDB1- dependent secreted factor involved in regulating proper muscle cell differentiation and fibrotic response.

## RESULTS

### Proteomic profiling reveals many secreted factors regulated by TGFβ and SETDB1 in DMD myotubes

Using human immortalized myoblasts derived from healthy individuals or DMD patients (Mamchaoui et al., 2011), we have previously shown that CM from TGFβ-treated myotubes impairs the receiving myoblasts differentiation and promotes the expression of pro-fibrotic genes (Granados et al., 2024). Interestingly, SETDB1 knockdown (KD) in the CM-producing myotubes partially counteracts this effect (Granados et al., 2024).

To identify the secreted factors mediating the inhibitory effect of the TGFβ-treated myotubes on the myoblast differentiation, we performed mass spectrometry (MS) analysis of the CM. Specifically, we analyzed CM collected from DMD myotubes upon SETDB1 KD and/or TGFβ treatment (**Fig 1A**). A total of almost 3000 proteins were identified across all samples (**Fig 1B, S1A,** and **Table S1**). The Principal Component Analysis (PCA) showed clear clustering of the samples, with the main separation driven by TGFβ treatment and SETDB1 KD (**Fig 1C**).

**Figure 1.**
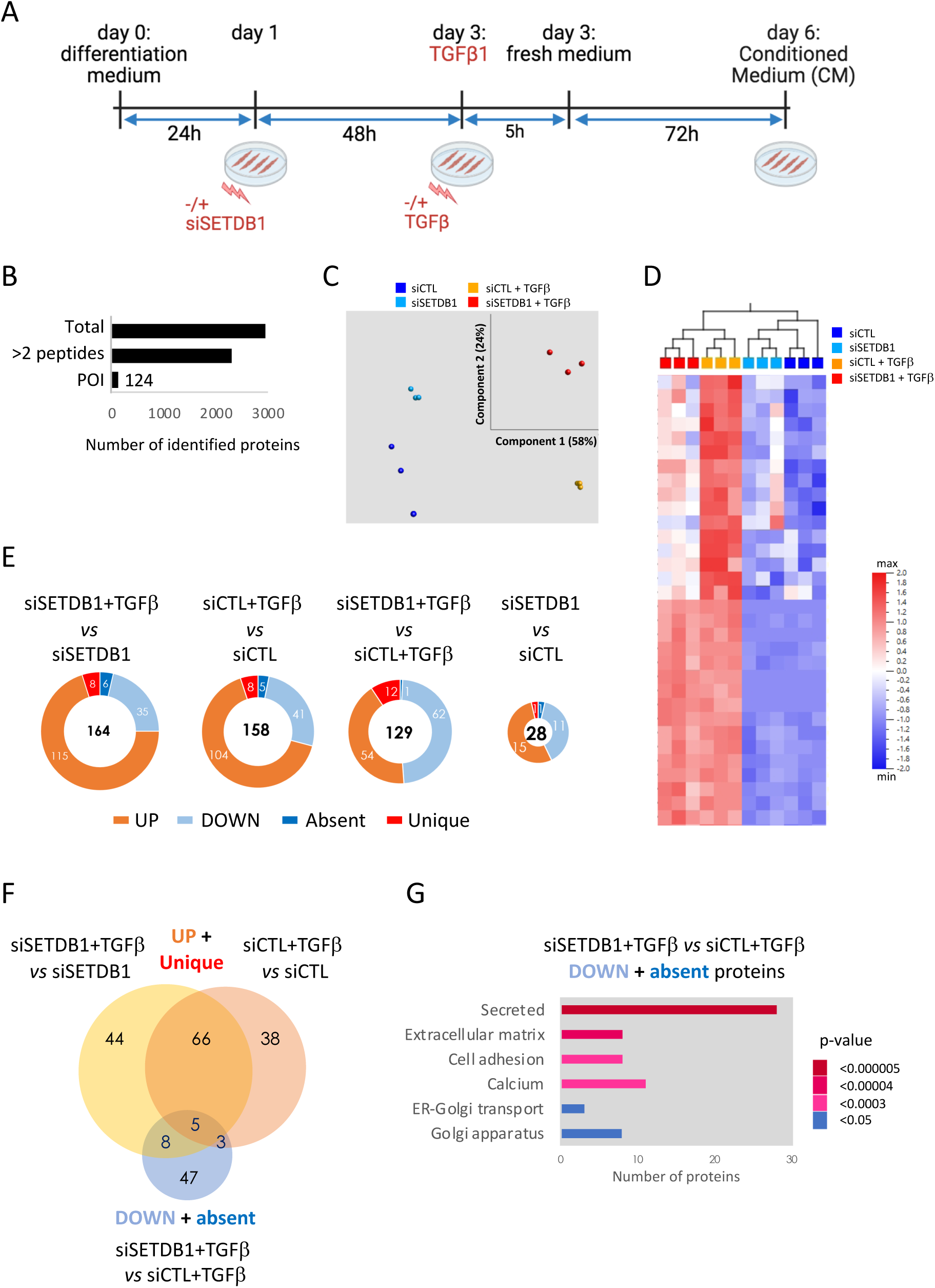
Proteomic profiling of the TGFβ-responsive and SETDB1-dependent secretome in DMD myotubes. **A.** Diagram of the secretome experiment. **B.** Secretome analysis of the mass spectrometry results on the condition media described in A (N=3, DMD #1), the complete list of identified proteins is shown in Table S1. Proteins of interest (POI) from the ANOVA analysis with p-value <0.001. **C.** Principal component analysis (PCA) of the significant target from the ANOVA analysis. **D.** Selected area of the heatmap (ANOVA. p-value < 0.001) showing the candidates that are upregulated upon TGFβ treatment (yellow) compared to basal condition (blue) and that decrease in siSETDB1+TGFβ (red). Complete heatmap is shown in Fig S1A. **E.** Doughnut plot showing the number of proteins from secretome analysis that are altered (down-regulated. up-regulated. absent or uniquely found). Total number of significant proteins is also indicated (T-test. p<0.01). A complete list of altered proteins along with the fold changes is shown in Table S2-S5. **F.** Venn diagram of the shared proteins between the two by two comparison from panel E. The Venn Diagram shows the intersection between up+unique proteins from comparison with siSETDB1+TGFβ vs siSETDB1 (impact of TGFβ in the SETDB1 KD) and with siCTL+TGFβ vs siCTL (impact of TGFβ) with the proteins down regulated and absent in siSETDB1+TGFβ vs siCTL+TGFβ (impact of SETDB1 KD on TGFβ treatment). **G.** David bioinformatics functional annotation chart (Sherman et al., 2022) using uniprotKB keywords (biological process, cellular component, molecular function and ligand) using down and absent protein from siSETDB1+TGFβ versus siCTL+TGFβ comparison.

Overall, we found a significant number of differentially enriched proteins in response to TGFβ treatment and/or SETDB1 KD (**Fig 1D-E, S1A**). We reasoned that relevant candidates would be upregulated in response to TGFβ, reflecting their potential to promote fibrosis and impair differentiation. Conversely, these factors should be downregulated upon SETDB1 KD, a condition that partially attenuates the pathological effects of TGFβ (Granados et al., 2024). Therefore, we focused on the proteins upregulated upon TGFβ treatment and decreased with SETDB1 KD (**Fig 1D**).

The comparison between TGFβ-treated condition with and without SETDB1 KD (siSETDB1-TGFβ vs siCTL+TGFβ) revealed a nearly equal number of downregulated (62+1) and upregulated (54+12) proteins (**Fig 1E**). The complete list of differentially detected proteins is shown in **Table S2-S5**. The signature of up-regulated and unique proteins induced by TGFβ overlapped by nearly 60% between the control and SETDB1 KD conditions (**Fig 1F**). Activation of the TGFβ pathway, independently of SETDB1 KD, led to an increase of secreted proteins related to the ECM (**Fig S1B-C**). The promising candidates which were downregulated upon SETDB1 KD in the presence of TGFβ belong to the secreted and ECM-related protein categories (**Fig 1G**). For instance, the ECM remodeling enzymes MMP14 and ADAMTS4 were upregulated in response to TGFβ, and downregulated upon SETDB1 KD (**Fig S1D**). A similar pattern was observed for the ECM component COL1A2 and the immune-related factor IL11 (**Fig S1D**). Regulation of MMP14 and IL11 was further confirmed at the transcript level in myotubes producing the CM, although the reduction in transcript levels induced by SETDB1 KD was less pronounced than the corresponding decrease observed at the protein level (**Fig S1E**).

Together, these findings reveal many TGFβ-responsive, SETDB1-dependent secreted factors in myotubes that may ultimately impact myoblast differentiation and the fibrotic response.

### Comparative proteomic reveals differences between the basal secretome of human DMD and healthy myotubes

After identifying deregulated secreted proteins upon stimulation, we next examined the basal secretome of DMD muscle cells to uncover intrinsic differences from healthy controls, independent of exogenous triggers. For this we performed a comparative proteomic analysis of the CM collected from human DMD and healthy immortalized myotubes under basal conditions after 3 days of differentiation **(Fig 2A)**. We identified a total of approximately 1600 secreted proteins across all samples **(Fig 2B)**. PCA revealed a clear separation between DMD and healthy samples **(Fig 2C)**, indicating a distinct secretome signature associated with the disease state independently of the *DYSTROPHIN* mutation. Comparative secretome analysis **(Fig 2D-E)** revealed that DMD myotubes are enriched in proteins involved in RNA processing (HNRNPR, SYNCRIP), vesicular trafficking (RAB11B, SEC23A), metabolic enzymes (PGK1, PKM), and cytoskeletal regulation (ARPC1A, CDC42), while structural ECM components (LAMA5) and signaling modulators (TGFβ2, ADAM10) are depleted. GO enrichment analysis highlighted overrepresentation of pathways related to amino acid biosynthesis, the renin-angiotensin system, beta-alanine metabolism, calcium reabsorption, and other metabolic and cellular processes **(Fig 2F)**.

**Figure 2.**
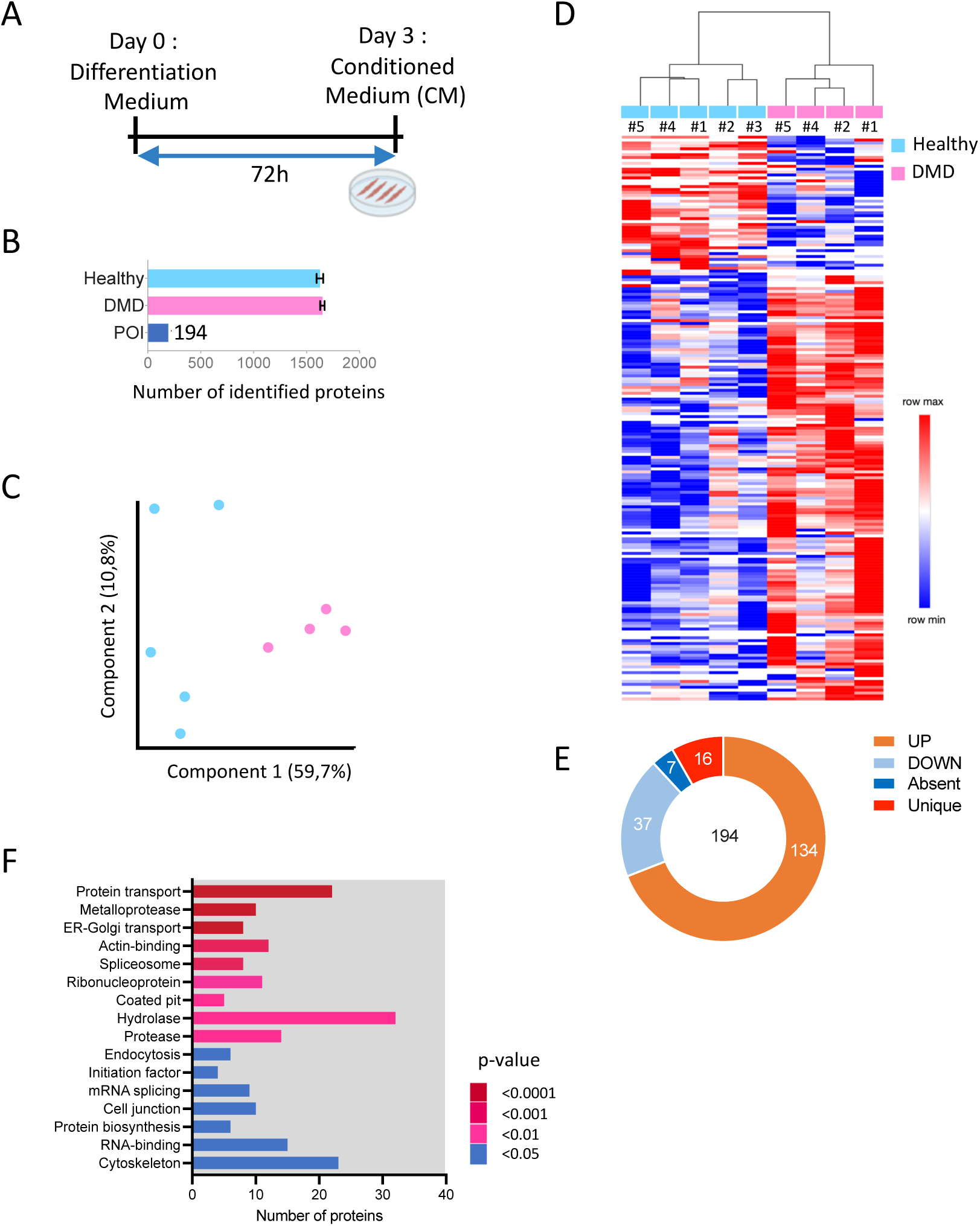
Proteomic profiling of basal secretome of DMD myotubes compared to healthy myotubes. **A.** Diagram of the secretome experiment. **B.** Secretome analysis of the mass spectrometry results on the conditioned media described in A (N=5 control and N=4 DMD). POI: proteins of interest. **C.** PCA of the significant target from the ANOVA analysis. **D.** Heatmap showing the POI (p-value < 0.05 and FC >1.2) differentially expressed in DMD samples (pink) compared to controls (blue). Detailed heat map is shown in Fig S2. **E** Doughnut plot showing the number of proteins differentially expressed in DMD samples compared to controls. **F.** Davidbioinformatics functional annotation chart (Sherman et al., 2022) using uniprotKB keywords (biological process, cellular component, molecular function and ligand) using all proteins of interest from the DMD secretome data.

Together, these results indicate a shift in the DMD secretome marked by enrichment of metabolic, RNA-binding, and cytoskeletal proteins, loss of structural ECM components and TGFβ signaling modulators, and the emergence of a DMD-specific secretory signature that may contribute to impaired myofiber stability and pathological remodeling of the muscle microenvironment.

### SETDB1 depletion reveals EMILIN1 as a key TGFβ-responsive ECM protein in DMD myotubes

In order to narrow down the list of altered proteins, we focused on the ones that are increased in DMD basal condition and also upregulated by TGFβ treatment but reduced with SETDB1 KD, by selecting proteins common to the DMD secretome and those downregulated (or absent) in the comparison siSETDB1+TGFβ *vs* siCTL+TGFβ. Our stringent analysis revealed only six candidates (EMILIN1, ARCN, NEXN, ADAM10, SSC5D, and NES), among them only EMILIN1 is upregulated in basal DMD condition, upregulated by TGFβ treatment and reduced with SETDB1 KD (**Fig 3A-B-C, S3A-B**). As a key ECM glycoprotein, EMILIN1 is involved in structural and regulatory functions. Indeed, EMILIN1 promotes cell adhesion and migration, and it acts as a negative regulator of TGFβ by blocking its maturation and activation (Colombatti et al., 2011). EMILIN1 protein level changes were confirmed also at the mRNA level, showing that DMD myotubes were indeed producing higher amounts of EMILIN1 upon TGFβ treatment and that SETDB1 KD partially reversed this effect (**Fig 3D**). DMD biopsies present high levels of fibrosis and variations in fiber size (Fig 3E) Interestingly, EMILIN1 was also found enriched, both at mRNA and protein levels in muscle tissue sections from DMD patients compared to healthy controls and localized around muscle fibers **(Fig 3E-F-G, Fig S3C).**

**Figure 3.**
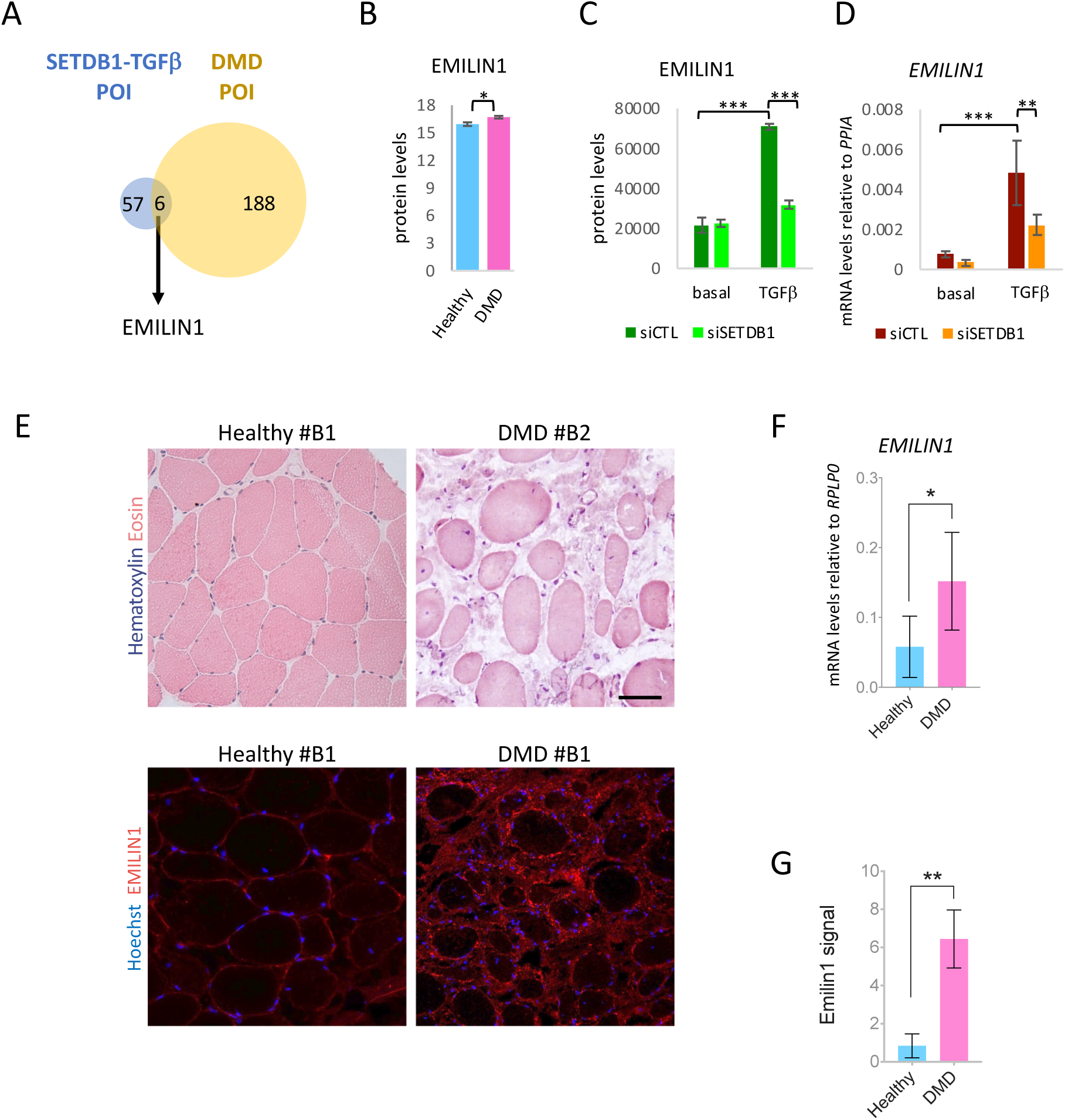
EMILIN1: Common target of the basal DMD and TGFβ–SETDB1 secretomes, upregulated in patient muscle. **A**. Venn Diagram of the shared proteins between the POI of DMD secretome (same as Fig 2E) and the downregulated proteins upon siSETDB1+TGFβ (same as Fig 1G). **B & C.** MS data analysis of EMILIN1 between healthy and DMD secretome (B) and DMD with SETDB1 KD +/- TGFβ (C). Data are represented as average +/-SEM *p<0.05; ***p<0.001 unpaired Student’s t test. **D.** RTqPCR of *EMILIN1* (N=3, DMD #1). Data are represented as average +/-SEM **p<0.01; ***p<0.001 unpaired Student’s t test. **E**. Hematoxylin and eosin coloration of healthy and DMD muscle biopsies– Immunostaining of EMILIN1 (red) and Hoechst (blue) scale bar = 50 µm. **F.** RTqPCR quantification of *EMILIN1* on human muscle biopsies of healthy and DMD patients (biological replicates N=7 Healthy-6 DMD). Data are represented as mean +/-SD *p<0.05 unpaired Student’s t test. **G**. Quantification of the percentage of EMILIN1 signal on healthy and DMD muscles (biological replicates N=3 Healthy N=4 DMD). Data are represented as mean +/-SD **p<0.01; unpaired Student’s t test.

Together, these findings identify EMILIN1 as a TGFβ and SETDB1-dependent target which is enriched in the DMD basal secretome and further upregulated in response to TGFβ stimulation. Notably, EMILIN1 overexpression in muscle biopsies from DMD patients supports its pathological relevance *in vivo*.

### EMILIN1 plays a role in myogenic differentiation of healthy and DMD myotubes

To better characterize the role of the ECM glycoprotein EMILIN1 in skeletal muscle cells, we studied its expression levels in healthy and DMD myoblasts during myogenic differentiation into myotubes (**Fig 4A**), monitoring this process with *MyoD* and *MyHC* (Myosin Heavy Chain) mRNA levels, which serve as canonical myogenic and differentiation markers (**Fig S4A**). Interestingly, EMILIN1 was more expressed in DMD proliferating myoblasts and myotubes at early stages compared to healthy controls (**Fig 4A**). This finding indicates that *EMILIN1* mRNA rises at the onset of differentiation, leading to the increased EMILIN1 protein secretion observed after 3 days of differentiation (**Fig 3B**).

**Figure 4.**
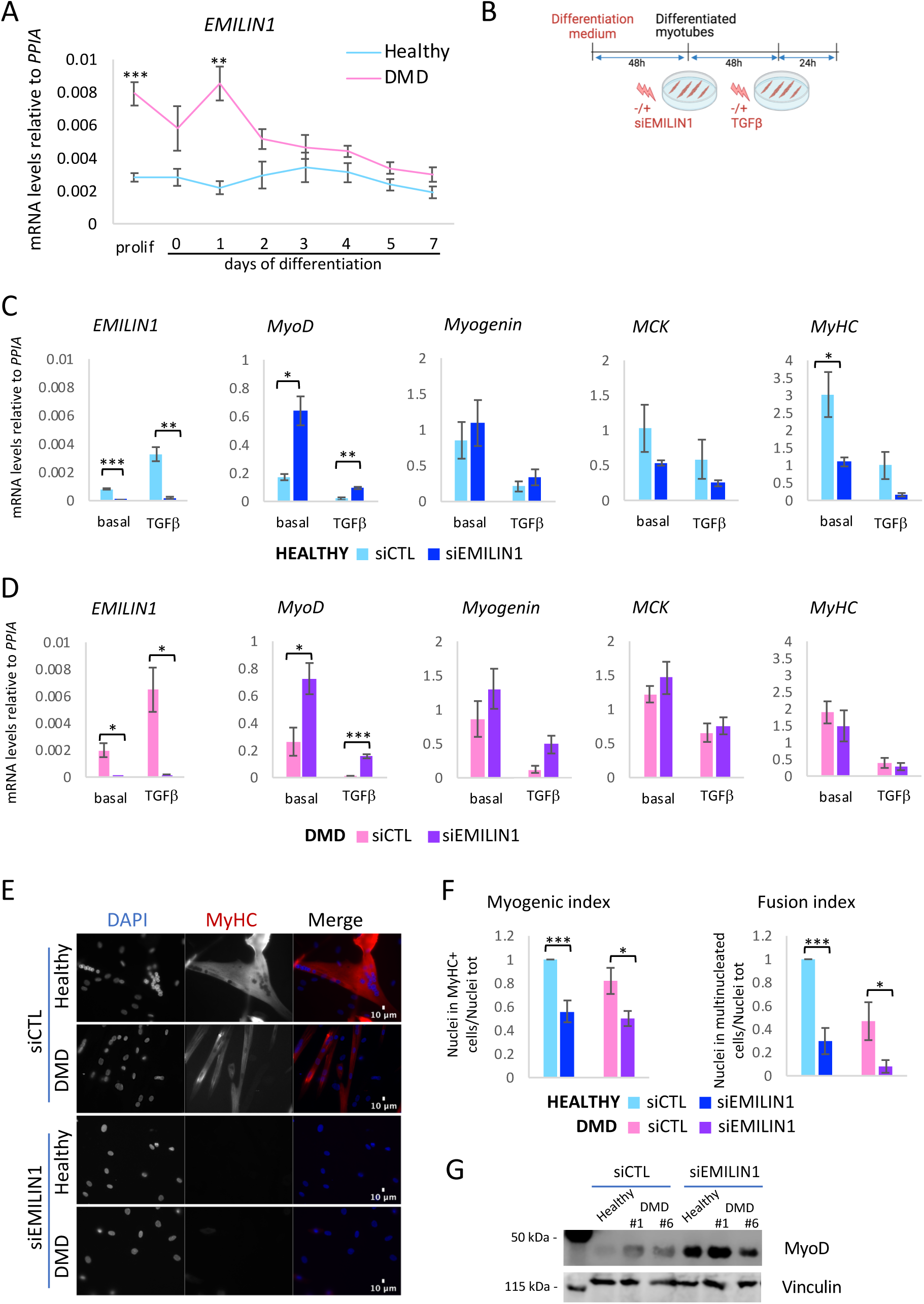
EMILIN1 KD increases the expression of early myogenic markers and inhibits late differentiation in healthy and DMD myotubes. **A.** RTqPCR of *EMILIN1* in healthy and DMD proliferating myoblasts and differentiated myotubes (N=4, DMD#1). **B.** Scheme of experimental design. Cells were transfected with siRNAs scrambled (siCTL) or against EMILIN1 (siEMILIN1) after 2 days of differentiation (myotubes) and 2 days later treated or not with TGFβ at 20ng/mL for 24h. **C-D.** RTqPCR of *EMILIN1* and of early (*MyoD* and *Myogenin*) and late (*MCK* and *MyHC*) myogenic markers in healthy (C) and DMD (D) myotubes with siCTL or siEMILIN1, +/- TGFβ (N=3, Healthy #1, DMD #1). **E.** Immunofluorescence of MyHC (red) and nuclei staining with DAPI (blue) in healthy and DMD myotubes with siCTL or siEMILIN1. Scale bar 10μm. A representative field is shown for each condition. **F.** Quantification of myogenic and fusion index (N=7, Healthy #1, DMD #1). For all panels, data are represented as average -/+ SEM *p<0.05; **p<0.01; ***p<0.001 unpaired Student’s t test. **G.** Western blot of the cells with siEMILIN1 or siCTL as in figure 4B, in absence of TGFβ treatment, showing protein levels of MyoD. Vinculin was used as loading control.

To further investigate the role EMILIN1 during myogenic differentiation, we performed siRNA-mediated KD experiments in proliferating myoblasts **(Fig S4B)** and differentiated myotubes (**Fig 4B**). The efficient KD of EMILIN1 in differentiated healthy and DMD myotubes led to a consistent and at least 2-times increase in the expression of the early master myogenic marker *MyoD*, both in basal condition and upon TGFβ treatment (**Fig 4C-D**). *Myogenin* expression, which acts downstream of MyoD during differentiation, was not consistently increased except when EMILIN1 KD was performed in proliferating myoblasts (**Fig S4C-S4D**). The expression of the intermediate myogenic marker *MCK* (muscle creatine kinase, a cytoplasmic enzyme involved in energy homeostasis) was only mildly impacted, showing almost no difference in DMD myotubes. However, the expression of the late myogenic marker *MyHC* was decreased upon EMILIN1 KD in differentiated healthy cells (**Fig 4C**) and both in healthy and DMD cells when EMILIN1 KD was performed in proliferating myoblasts (**Fig S4C-D**). The negative impact of EMILIN1 KD on the expression of *MyHC* was confirmed by the presence of less myotubes upon EMILIN1 depletion (**Fig 4E**), as represented by the quantification of the myogenic and fusion index (**Fig 4F**). To exclude that these effects were DMD mutation-dependent, we performed the same experiments on myotubes carrying a different mutation in the *DYSTROPHIN* gene (DMD #6). The efficient KD of EMILIN1 led to an increase of the expression of MyoD, not only at the mRNA (**Fig S4E-F**) but also at the protein level (**Fig 4G**), without a clear impact on *Myogenin*, while it reduced the expression of *MCK* and *MyHC* and results are similar with or without TGFβ (**Fig S4E-F**). The inability of the cells to reach the late stages of differentiation was confirmed by a decrease in the number of multinucleated cells (**Fig S4G**), as quantified by myogenic and fusion index (**Fig S4H**).

These results reveal a role of EMILIN1 in skeletal muscle differentiation, showing that its loss leads to increased expression of early myogenic marker *MyoD* but reduced expression of the late marker *MyHC*. This reduction results in fewer multinucleated myotubes, indicating impaired late-stage differentiation. These effects are consistent across both healthy and DMD myotubes, suggesting that controlled levels of EMILIN1 are important for proper muscle cell differentiation.

### EMILIN1 regulates the expression of the fibrotic marker *SERPINE1*

To assess the role of EMILIN1 in the TGFβ-mediated fibrotic response, we focused on *SERPINE1*, a fibrotic marker we have already studied in our previous work (Granados et al., 2024). PAI-1 (plasminogen activator inhibitor 1), the protein encoded by *SERPINE1*, was increased in the secretome of TGFβ-treated DMD myotubes, and it decreased upon SETDB1 KD (**Fig 5A**). Moreover, it was enriched in the secretome of DMD myotubes compared to healthy ones, being already detected in basal condition and further increased by TGFβ (**Fig 5B-S5A**). Remarkably, *SERPINE1* expression was impacted by EMILIN1. EMILIN1 KD in differentiated healthy and DMD myotubes induced a decrease in *SERPINE1* expression both in basal condition and upon TGFβ treatment (**Fig 5C**). This decreasing trend in *SERPINE1* mRNA levels was validated also upon EMILIN1 KD in proliferating myoblasts (**Fig S5B**) and in myotubes healthy #4 and DMD #6 (**Fig 5D**). Notably, the levels of PAI-1, were reduced following EMILIN1 KD, mirroring the changes observed at the mRNA level (**Fig 5E-F**). This further supports the observation that reducing EMILIN1 leads to a decrease in the fibrotic marker PAI-1.

**Figure 5.**
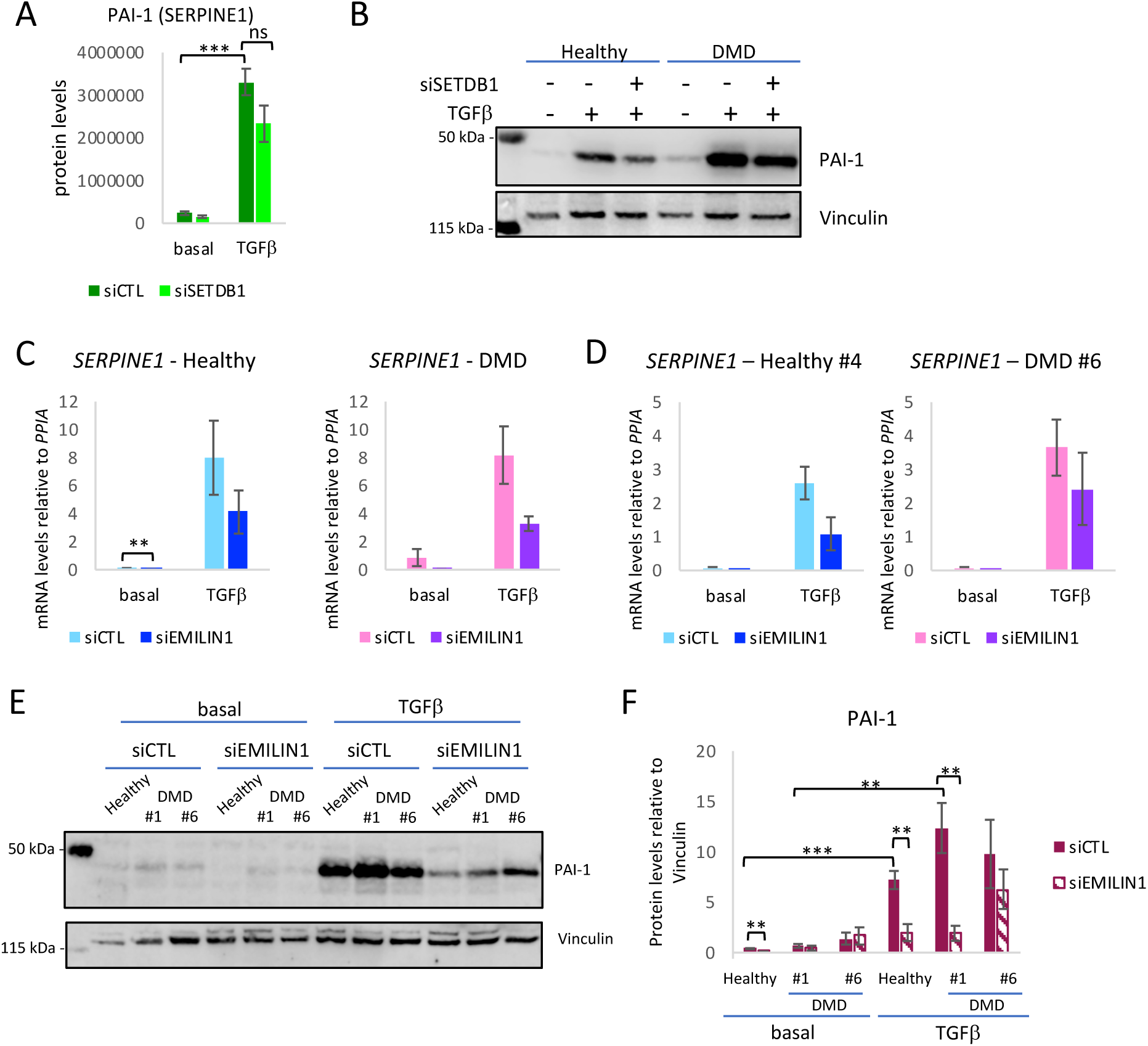
EMILIN1 KD decreases the expression of the TGFβ- and SETDB1- dependent fibrotic marker SERPINE1 in healthy and DMD myotubes. **A.** MS data analysis of PAI-1 (*SERPINE1*) **B.** Western blot of the CM obtained as in Figure 1A from healthy and DMD myotubes +/- siSETDB1, +/- TGFβ showing protein levels of PAI-1. Vinculin was used as loading control (N=3, Healthy #1, DMD #1). Quantification is shown in Fig S5A. **C-D.** RTqPCR of *SERPINE1* in healthy and DMD myotubes as in figure 4B (N=3, Healthy #1, DMD #1 for C and Healthy #4, DMD #6 for D). **E.** Western blot of the myotubes treated with siEMILIN1 and TGFβ as in figure 4B showing protein levels of PAI-1. Vinculin was used as loading control. **F.** Quantification of the western blot 5E normalized on Vinculin (N=4, Healthy #1, DMD #1 and DMD #6). For all panels, data are represented as average -/+ SEM *p<0.05; **p<0.01; ***p<0.001 unpaired Student’s t test.

Together, these findings suggest that EMILIN1 plays a role in regulating the TGFβ-mediated fibrotic response, as indicated by the modulation of the fibrotic marker SERPINE1.

## DISCUSSION

The regenerative capacity of skeletal muscle depends on the dynamic interplay between its various components, including muscle fibers and the surrounding connective tissue. This tissue operates as an integrated system, finely tuned to respond to injury and maintain muscle function (Mukund & Subramaniam, 2020). While skeletal muscle is primarily known for its role in contraction, it also functions as an active endocrine organ, capable of releasing a variety of signaling factors. These factors enable muscle cells to communicate in an auto-, para-, or endocrine manner, influencing various physiological processes, including muscle regeneration (Florin et al., 2020). Muscle regeneration is a highly coordinated process, that depends on the precise secretion of multiple factors to orchestrate tissue repair. Through the complex communication between muscle cells, immune cells and Fibro-Adipogenic Progenitors (FAPs), skeletal muscle can effectively repair itself and restore functionality. Notably, in DMD, muscle regenerating abilities are compromised. The sustained inflammation leads to TGFβ-induced fibrosis, defined as an excessive accumulation of ECM components, and to fat accumulation. Over time, functional muscle tissue is replaced with non-functional fibrotic and fatty tissue, resulting in progressive muscle loss and fibro-fatty tissue accumulation (Serrano et al., 2011).

Our previous work has established that the conditioned medium from TGFβ-treated myotubes has a negative impact on myoblast differentiation and promotes fibrosis. SETDB1 KD is able to partially counteract this effect, indicating that SETDB1 potentiates the TGFβ-induced fibrosis in DMD muscles. However, even though targeting SETDB1 in postmitotic myotubes and in FAPs does not seem to have a deleterious effect on cell viability, its targeting in muscle tissue would be challenging, since the impact on other cell types, such as macrophages, or on the crosstalk between cell types, is unknown (Garcia et al., 2024; Granados et al., 2024). Therefore, it appears crucial to identify the SETDB1/TGFβ-dependent secreted factors which could play a role in the deleterious environment of DMD.

Our data presented here describe the effect of SETDB1 regulation on the TGFβ pathway, in particular regarding the composition of the DMD secretome. The extensive mass spectrometry analysis revealed the presence of almost 3000 proteins in the secretome of DMD myotubes. The good clustering of the samples highlights that SETDB1 and TGFβ are indeed impacting the identity of the secretome. To better understand the effect of the CM described in our previous work (Granados et al., 2024), we focused on proteins that are upregulated by TGFβ, potentially contributing to impaired myoblast differentiation and enhanced fibrosis, but are downregulated upon SETDB1 KD, where the described detrimental effects are attenuated.

In addition to the changes induced by TGFβ and SETDB1, our data also provide insights into the composition of the basal DMD secretome. Notably, several secreted factors, including ECM and signaling proteins, are already present at significant levels in untreated DMD myotubes. This suggests that even in the absence of exogenous TGFβ stimulation, the DMD muscle environment is primed toward a fibrotic and regenerative-impairing profile. Interestingly, the integrated results of both secretome analyses reveal EMILIN1 among the secreted proteins. This ECM glycoprotein is not only upregulated in response to TGFβ and downregulated upon SETDB1 knockdown, but also present in the basal DMD secretome, where it shows higher levels compared to healthy controls. Moreover, EMILIN1 is overexpressed in patient biopsies, suggesting a dual mode of regulation and a sustained involvement in the pathological environment.

EMILIN1 is a protein involved in elastic and collagen fibers formation, whose role in skeletal muscle remains largely unexplored. Interestingly, EMILIN1 possesses a gC1q domain, which mediates interactions with integrins, contributing to cell adhesion and migration, and an EMI domain with seven cysteine residues, involved in TGFβ processing (Colombatti et al., 2011). EMILIN1 is described as an inhibitor of the TGFβ pathway in different contexts, such as cancer and skin psoriasis (Honda et al., 2024; Pivetta et al., 2022; Zacchigna et al., 2006). Although *EMILIN1* knockout mice appear grossly normal, they manifest specific phenotypes including skin defects and hypertension (Zanetti et al., 2004). In humans, two distinct missense mutations within the *EMILIN1* gene have been identified: c.64G>A (p.A22T) in exon 1, associated with connective tissue disorder and sensory-motor neuropathy; and c.748C>T (p.R250C) in exon 4, linked to distal motor neuropathy (Capuano et al., 2016; Iacomino et al., 2020). However, EMILIN1 role in skeletal muscle and, specifically in DMD, is unknown.

The detection of EMILIN1 under basal conditions further supports its relevance as a constitutive component of the dystrophic secretome, whose function may be altered independently of acute TGFβ modulation. Importantly, EMILIN1 is not only detected in the basal secretome of DMD myotubes, but is also found to be upregulated in muscle tissue sections from DMD patients, reinforcing its relevance *in vivo* and suggesting that its role in DMD pathology extends beyond *in vitro* models.

Here, we show that EMILIN1 KD in myoblasts and differentiated myotubes has an impact on the differentiation process. Specifically, EMILIN1 KD led to an increase in the early myogenic marker MyoD. This result is in accordance with experiments done in *Xenopus* where overexpression of EMILIN1 or only EMI-domain in eight cell stage embryos reduced MyoD expression and inhibited mesoderm development (Zacchigna et al., 2006). Despite MyoD upregulation, we observe a decrease in the late myogenic marker MyHC. The reduction in multinucleated myotubes upon EMILIN1 KD further highlights the role of EMILIN1 in promoting proper muscle fiber maturation. Notably, the same pattern of differentiation impairment was observed in both healthy and DMD myotubes, suggesting that EMILIN1 regulates muscle differentiation in a similar manner regardless the presence of *DYSTROPHIN* mutations. Whether EMILIN1 regulates myogenic differentiation through the TGFβ pathway, which is known to prevent myoblasts fusion (Girardi et al., 2021), or through alternative mechanisms is yet to be explored.

Additionally, we show that EMILIN1 KD leads to a reduction in the expression of the fibrotic and TGFβ-dependent marker *SERPINE1*. *SERPINE1* codes for the plasminogen activator inhibitor 1 (PAI-1), a protein localized in the extracellular space which plays critical roles in response to skeletal muscle injury and in myopathies. PAI-1 contributes to the inhibition of ECM degradation, and excessive PAI-1 level promotes fibrosis and results in impaired skeletal muscle regeneration (Rahman & Krause, 2020).

Given the inhibitory role of EMILIN1 on TGFβ signaling in other contexts, it may seem counterintuitive that its reduction would lead to decreased fibrosis, potentially improving the DMD environment. However, this finding might be context-dependent and further studies are needed to elucidate the interplay between EMILIN1 and the TGFβ pathway in DMD. In particular, studying the role of EMILIN1 in FAPs, key mediators of the fibrotic response, could provide valuable insights into how this secreted factor contributes to the DMD phenotype.

In summary, these findings reveal critical insights into the interplay between SETDB1, TGFβ signaling, and the fibrotic response in DMD. The data highlight the role of SETDB1 in modulating the secretome of DMD myotubes and provide a rich list of factors that could contribute to the pathological environment characteristic of DMD. Importantly, our data show that several components of this secretome, including EMILIN1, are already present in the basal dystrophic state and are upregulated in patient muscle sections, suggesting a pre-existing and persistent alteration of the secretory profile in DMD muscle. Among these factors, EMILIN1 emerged as an important player involved in the regulation of myogenic differentiation and fibrosis. While EMILIN1 targeting appears promising for reducing fibrosis, it does not enhance terminal myotube differentiation, indicating that combinatorial approaches may be necessary to fully restore muscle function. Collectively, our results suggest that SETDB1-dependent secreted factors can influence both myoblast differentiation and fibrotic signaling, offering novel potential targets for therapeutic intervention in DMD.

## MATERIALS AND METHODS

### Human muscle samples

Human muscle samples from healthy controls and Duchenne muscular dystrophy patients (Table 1) were obtained via the Myobank-AFM, affiliated with EurobioBank, in accordance with European recommendations and French legislation (authorization AC-2019-3502).

**Table 1.**
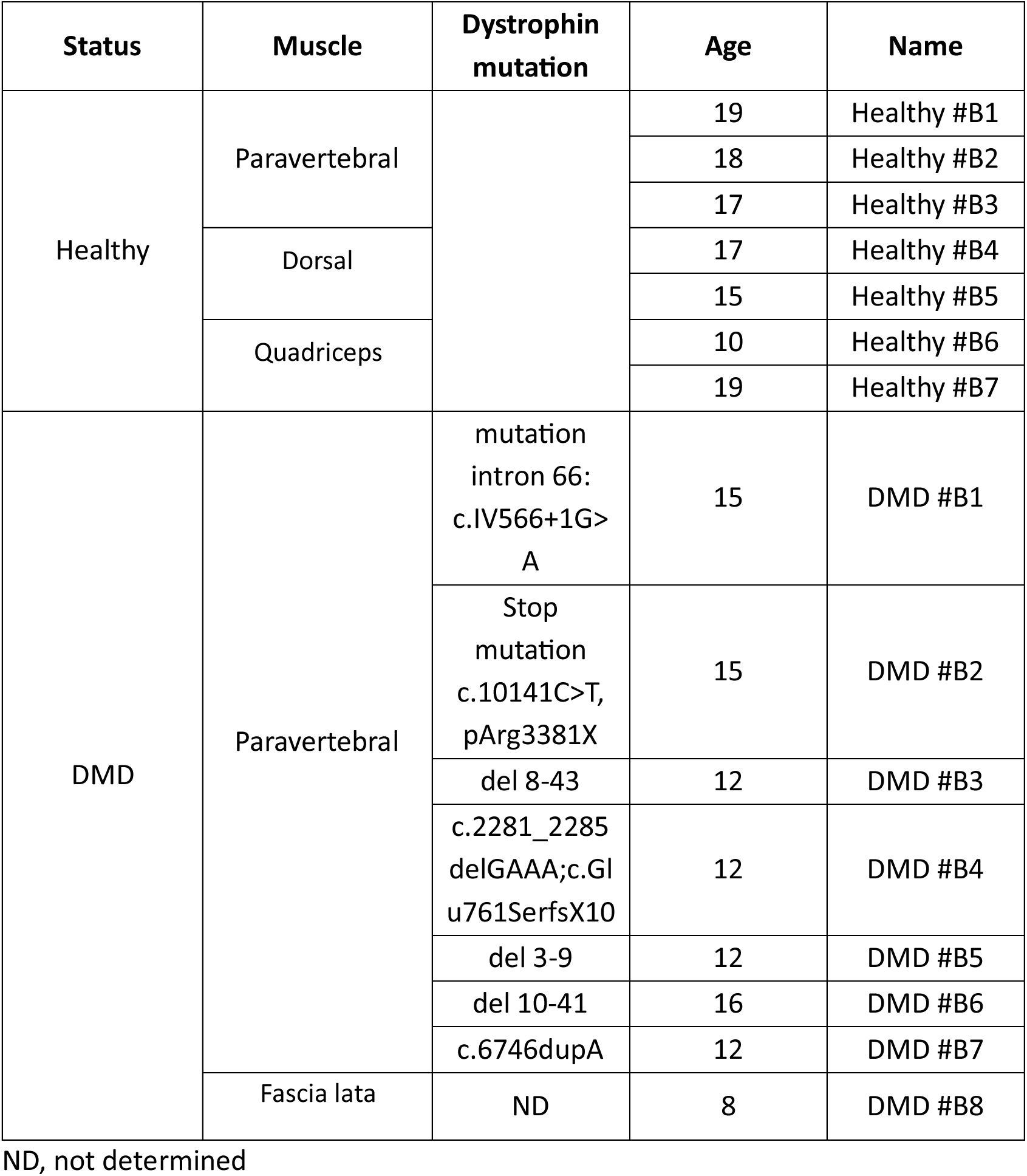
Muscle biopsies.

### Cell culture

Human control and DMD immortalized myoblasts were obtained from the MyoLine platform for immortalization of human cells from the Institute of Myology (Paris, France) (Table 2). Immortalization was performed as described in (Mamchaoui et al., 2011), using human Telomerase-expressing (hTERT) and Cyclin-Dependent Kinase 4 (CDK4)–expressing vectors. They were cultured on gelatin-coated plates for all the experiments except for the preparation of the secretome for the healthy *versus* DMD comparison. Immortalized myoblasts were cultured in growth medium (GM) composed of Dulbecco’s modified Eagle’s medium (DMEM) high glucose GlutaMAX (Invitrogen, 61965-026)/Medium 199 (Invitrogen, 41150020) 4:1 mixture, supplemented with 20% fetal bovine serum (Dutscher, 500105M1M batch S00G0), Fetuin (25 μg/ml; Sigma-Aldrich, F3385), Insulin (5 μg/ml; Sigma-Aldrich, I9278), basic Fibroblast Growth Factor (0.5 ng/ml; Sigma-Aldrich, SRP3043), human Epidermal Growth Factor (5 ng/ml; Sigma-Aldrich, SRP3027), and dexamethasone (0.2 μg/ml; Sigma-Aldrich, D4902).

**Table 2.** Cell lines.

| Status | Muscle | Dystrophin mutation | Age | Name |
| --- | --- | --- | --- | --- |
| Healthy | Paravertebral |  | 16 | Healthy #1 |
|  |  |  | 13 | Healthy #2 |
|  |  |  | 12 | Healthy #3 |
|  | Fascia lata tensor |  | 20 | Healthy #4 |
|  | Gracilis |  | 25 | Healthy #5 |
| DMD | Paravertebral | Del 45-52 | 13 | DMD #1 |
|  |  | Del 51-54 | 15 | DMD #2 |
|  |  | Mutation intron 70:<br>C.IVS70+1G>A | 17 | DMD #3 |
|  | Spinal | Del 45 | 14 | DMD #4 |
|  | Dorsal | Duplication exon 10 to 11 | 14 | DMD #5 |
|  | Quadriceps | stop mutation exon 59:<br>c.8713C>T,<br>p.Arg2905X | 11 | DMD #6 |

### Myogenic differentiation of human immortalized myoblasts

Human control and DMD-immortalized myoblasts were grown at >80% confluence, and the medium was switched for DM media. The DM media was composed of DMEM with gentamycin for the basal DMD vs control secretome comparison and of DMEM with insulin (10μg/mL; Sigma-Aldrich, I9278) for all the other experiments, as described in (Granados et al., 2024).

### Myoblast and Myotube transfection with siRNAs

Proliferating myoblasts (cultured in GM) or two-day-differentiated myotubes (cultured in DM) were transfected with siRNAs at a final concentration of 70nM, using the Lipofectamin RNAiMAX transfection agent (Invitrogen, 13778100). Cells were kept for 2 days in a transfection medium before switching to fresh media. We used the ON-TARGETplus Human SETDB1 siRNA (Dharmacon, L-020070-00-0010); EMILIN1 siRNA (Sigma-Aldrich, SASI_Hs01_00218611), and the scrambled control siRNA (Sigma-Aldrich, F: CAUGUCAUGUGUCACAUCUC[dT][dT], R: GAGAUGUGACACAUGACAUG[dT][dT]).

### Immunofluorescence

Human myoblasts were grown on gelatin-coated coverslips. They were fixed with 4% paraformaldehyde for 20 min, and they were saturated with 50 mM NH4Cl for 10 min at room temperature (RT). Then, they were permeabilized with 0.5% Triton X-100 for 5 min. Cells were washed three times with PBS between each step. Cells were next incubated with primary antibodies at the concentration indicated in the antibody list below. Alexa Fluor 555 was used as secondary antibodies (Invitrogen, 115-165-207). Nuclei were counterstained with 4ʹ,6-diamidino-2-phenylindole (DAPI). Cells were washed three times with PBS-Tween 0.1% FBS 2% after each antibody incubation.

Images were acquired with a Leica DMI-6000B microscope and analysis was performed using Fiji software.

For immunostaining of muscle biopsies, cryosections were fixed in 4% paraformaldehyde for 10 min at RT and incubated with PBS containing 2% FBS and 0,5% triton for 30 min at RT. Incubation with primary antibodies was performed at RT for 1 hour and cryosections were incubated with appropriate fluorescent secondary antibodies (Life technologies) at 1/400 at RT for 45 min. Nuclei were stained with Hoechst. Widefield fluorescence images were taken with an Axio Observer 7 microscope (Zeiss) equipped with a motorized stage coupled to an Orca Flash 4 Camera (Hamamatsu) and driven by the Zen software (Zeiss). The antibodies used are listed in Table 3.

**Table 3.**
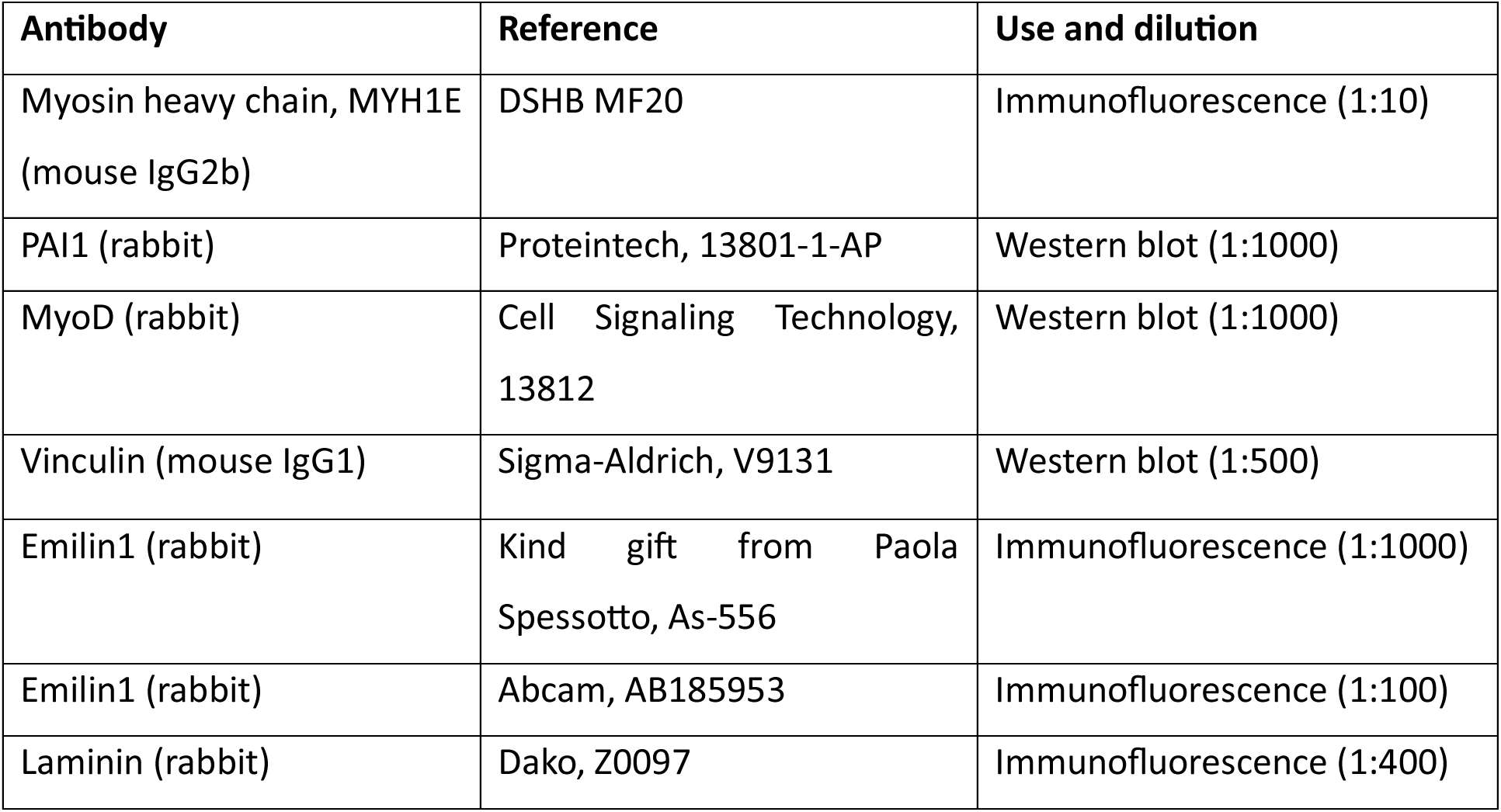
Antibodies.

| Antibody | Reference | Use and dilution |
| --- | --- | --- |
| Myosin heavy chain, MYH1E (mouse IgG2b) | DSHB MF20 | Immunofluorescence (1:10) |
| PAI1 (rabbit) | Proteintech, 13801-1-AP | Western blot (1:1000) |
| MyoD (rabbit) | Cell Signaling Technology, 13812 | Western blot (1:1000) |
| Vinculin (mouse IgG1) | Sigma-Aldrich, V9131 | Western blot (1:500) |
| Emilin1 (rabbit) | Kind gift from Paola Spessotto, As-556 | Immunofluorescence (1:1000) |
| Emilin1 (rabbit) | Abcam, AB185953 | Immunofluorescence (1:100) |
| Laminin (rabbit) | Dako, Z0097 | Immunofluorescence (1:400) |

### RNA extraction and reverse transcription quantitative PCR (RTqPCR)

Total RNA from cells was extracted using RNeasy Micro Kit (Qiagen, 74004) following the manufacturer’s procedures. Deoxyribonuclease (Qiagen, 79254) treatment was performed to remove residual DNA. One microgram of total RNA was reverse-transcribed with a High-Capacity cDNA Reverse Transcription Kit (Applied Biosystems, 4368813). Real-time quantitative PCR was performed to analyze relative gene expression levels using SYBR Green GoTaq® qPCR Master Mix (Promega, A6002) following the manufacturer’s indications. Relative expression values were normalized to the housekeeping gene mRNA *PPIA*.

For human frozen muscle sections, RNA was extracted using Trizol reagent (Invitrogen) according to the manufacturer’s instructions. RNA was reverse transcribed using M-MLV (Invitrogen) according to the manufacturer’s instructions. Quantitative polymerase chain reaction (qPCR) was carried out using SYBR Green Mastermix (Roche Applied Science) in a Applied Biosystems QuantStudio 6 Pro (Thermofisher scientific) with the following cycling protocol: 8 min at 95°C; followed by 50 cycles at 95°C for 15 s (s), 60°C for 15 s and 72°C for 15 s, and a final step consisting of 5 s at 95°C and 1 min at 65°C. The specificity of the PCR product was evaluated by melting curve analysis using the following program: 65°C to 97°C with a 0.11°C/s increase and gene expression levels were normalized to human RPLP0 and quantified with the 2–ΔΔCt method. Primers are listed in Table 4.

**Table 4.**
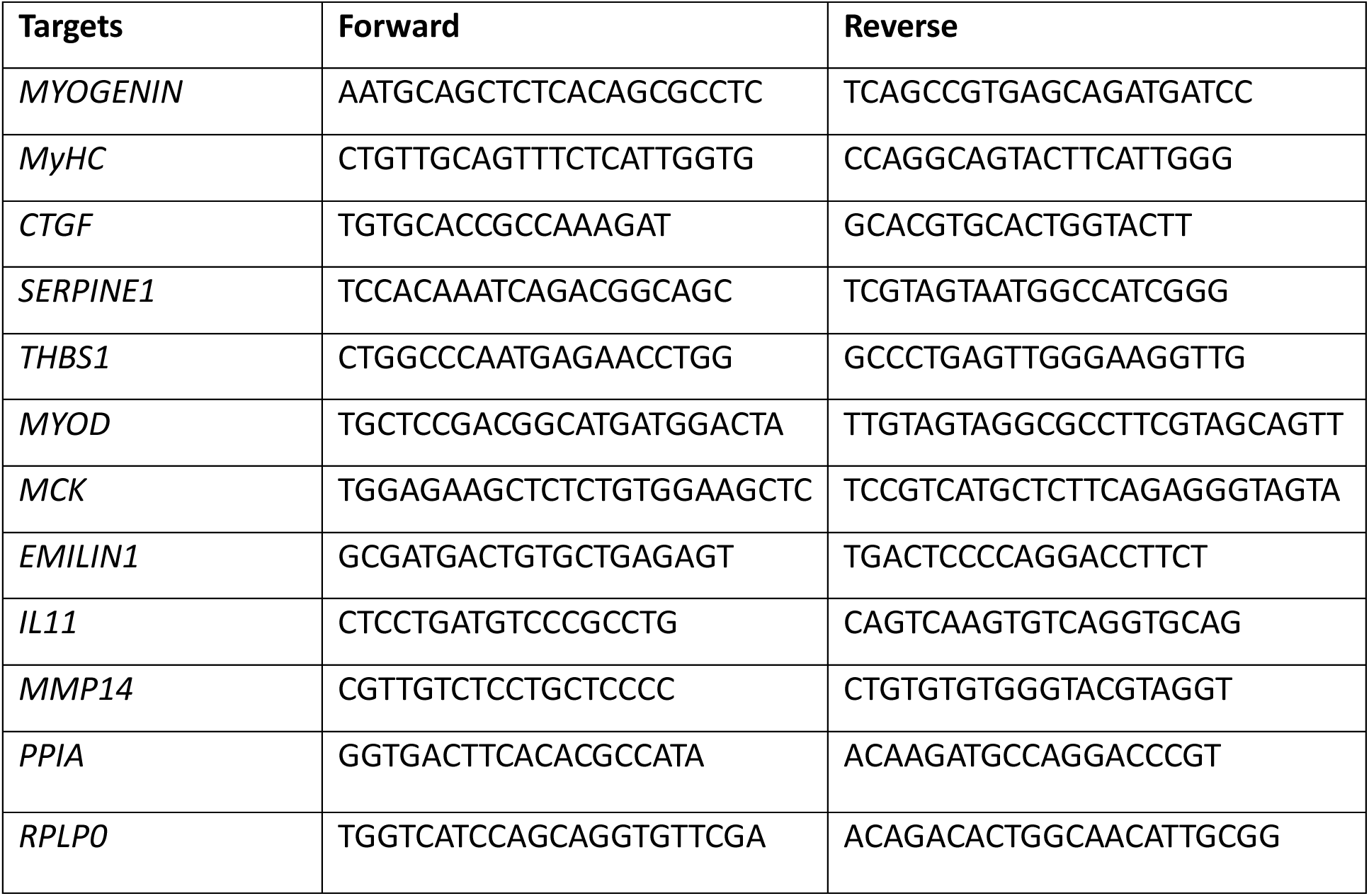
Primers.

### Western blot

Cells were lysed in RIPA buffer [20 mM tris (pH 7.65), 150 mM NaCl, 0.1% SDS, 0.25% NaDoc, and 1% NP-40] supplemented with protease inhibitor 1× (Sigma-Aldrich, SIGMAFAST S8830) and phosphatase inhibitor 1× (Sigma-Aldrich, P5726-5ML). Cell lysates were sonicated at 4°C for 10 min (30 s ON, 30 s OFF) at medium frequency (Bioruptor Diagenode). Then, the lysates were centrifugated for 10 min at 4°C at 10000 rcf and the supernatants were kept as the samples. Extracts were resolved on pre-cast polyacrylamide gel cassettes (NuPAGE 4 to 12% Bis-Tris) (Invitrogen NP0335BOX, NP0336BOX) and 1× NuPAGE MOPS SDS Running Buffer (Invitrogen, NP0001) and transferred into nitrocellulose membrane (Sigma-Aldrich, Amersham, 10600002) in 20 mM phosphate transfer buffer (pH 6.7). The membrane was blocked in 5% skim milk in PBST buffer (1× PBS, 0.2% Tween 20) and incubated overnight at 4°C with the primary antibody (Table 3). Membranes were washed twice for 5 min in PBST, incubated with appropriate secondary antibody IRDye (Li-Cor, IRDye 800CW, IRDye 680RD) in PBST, washed twice for 10 min in PBST, once for 10 min in PBS, and then imaged on Odyssey Imaging System.

### Collection and Preparation of Conditioned Media for WB and Analysis by Mass Spectrometry (MS)

Human myoblasts were cultured and differentiated as described above (serum-free DM). On day 1 of differentiation, cells were transfected with siSETDB1. Two days post transfection, cells were treated with TGFβ (20ng/mL; Peprotech, 100-21C-10UG) for 5 hours. Then, cells were washed five times with PBS before putting fresh DM. The CM was collected 72h later and filtered using a 0.2 μm filter to ensure removal of any dead cells. The CM was concentrated using spin columns with 10kDa cut off (Millipore Amicon Ultra, UFC5010) performing a centrifugation of 10 min at 10000 RCF, 4°C. The protein content of the CM was measured with BCA assay (ThermoFisher, Pierce BCA Protein Assay Kit, 23227). The concentrated CM was analyzed by WB (as described above) or by MS (10 μg of proteins per condition were analyzed as follows).

### Mass spectrometry

Mass spectrometry LC-MS/MS analyses were performed on two sites: at Proteom’IC, the proteomics core facility of Institut Cochin in Paris (https://institutcochin.fr/plateformes/proteomic) for the basal DMD vs control comparison and at ProteoSeine, the proteomics facility of Institut Jacques Monod in Paris (https://www.ijm.fr/plateformes-et-plateaux-techniques/proteoseine/) for the TGFβ and SETDB1 study. The MS proteomics data have been deposited to the ProteomeXchange Consortium via the PRIDE partner repository with the dataset identifier PXDXXXXX.

#### Samples preparation prior to LC-MS/MS analysis

Protein samples from both protocols underwent denaturation, reduction, alkylation, enzymatic digestion and peptide purification prior to LC-MS/MS analysis. However, specific conditions were adapted to the biological context and experimental constraints.

For the DMD secretome, proteins were extracted in a lysis buffer containing 400 mM triethylammonium bicarbonate, 4% SDS, 100 mM chloroacetamide, and 20 mM tris(2- carboxyethyl)phosphine, followed by heating at 95 °C for 5 minutes. Protein digestion was performed using trypsin (Promega) in S-Trap Micro Spin Columns (Protifi), following the manufacturer’s instructions. The resulting peptides were dried by SpeedVac and resuspended in 10 µL of 0.1% trifluoroacetic acid (TFA) with 10% acetonitrile (ACN). In cases of column clogging, additional clean-up was performed using strong cation exchange (SCX) membranes (3M Empore) (Rappsilber et al., 2007). Peptides were resuspended in 2% TFA, passed through SCX disks packed in modified P200 tips, washed with TFA-based buffers, and eluted with 5% ammonium hydroxide in 80% ACN. The eluates were dried and reconstituted in the same loading buffer as above.

For the SETDB1-TGFβ secretome, a six-time volume of cold acetone (−20°C) was added to a sample volume containing 10 µg of protein extracts. Vortexed tubes were incubated overnight at -20°C then centrifuged for 10 min at 11000 rpm, 4°C. The protein pellets were dissolved in urea 8M - NH4HCO3 25mM buffer. Samples were then reduced with DTT 10 mM and alkylated with IAA 20 mM. After a 16-fold dilution in NH4HCO3, samples were then digested overnight at 37°C by a mixture of trypsin/Lys C (1/20 Enzyme/Substrate ratio). The digested peptides were loaded and desalted on Evotips provided by Evosep one (Odense, Denmark) according to manufacturer’s procedure.

#### LC-MS/MS acquisition

Peptides from both protocols were analyzed using a timsTOF Pro 2 mass spectrometer (Bruker Daltonics, Bremen, Germany), operated in data-dependent acquisition with Parallel Accumulation-Serial Fragmentation (PASEF) mode (Meier et al., 2015), and coupled either to a Dionex U3000 RSLC nano-LC system (DMD samples) or to an Evosep one system (Evosep, Odense, Denmark) operating with the 30SPD method developed by the manufacturer (SETDB1-TGFβ samples). Chromatographic separation was performed on C18 reverse-phase columns under nanoflow conditions, using a binary solvent system consisting of water with 0.1% formic acid (solvent A) and acetonitrile with 0.1% formic acid (solvent B). For the SETDB1-TGFβ samples, peptides were separated using a 0.15 × 150 mm C18 analytical column (1.9 µm particles, EV-1106; Evosep) operated at 40 °C and 500 nL/min under the manufacturer’s 30 samples-per-day method. A 44-minute linear gradient was applied, resulting in a total run time of 48 minutes. In contrast, DMD and healthy samples were analyzed using an Aurora C18 column (75 µm × 25 cm, 1.7 µm particles, IonOpticks) over a 2-hour gradient, ranging from 98% solvent A to 35% solvent B. For SCX-cleaned samples, a shorter multistep gradient was used (10% B in 14 min, 20% B in 25 min, 40% B in 38 min). In all cases, the timsTOF instrument operated in positive ion mode, with precursor scans acquired in the 100–1700 m/z range. Ion mobility spectrometry (TIMS) was enabled, with accumulation and ramp times set between 166 ms and 180 ms. The ion mobility range was set to 1/K₀ = 0.6–1.6 Vs/cm² (DMD) or 0.75–1.25 Vs/cm² (SETDB1-TGFβ). PASEF cycles included 6 (SETDB1-TGFβ) or up to 10–12 MS/MS scans (DMD), achieving near-complete duty cycles.

Precursor ions were selected based on intensity thresholds (>1000 a.u.) and dynamically excluded for 0.4 s (DMD) or 0.8 min (SETDB1-TGFβ). They were re-sequenced until reaching a target intensity of 20,000 a.u. for DMD or 16,000 a.u. for SETDB1-TGFβ. Quadrupole isolation was synchronized with TIMS elution profiles, and collision energy was ramped based on ion mobility in all cases.

#### MS data analysis

Data from the DMD secretome were processed using MaxQuant (version 2.0.3.0) ) (Tyanova et al., 2015). Spectra were searched against a database combining SwissProt human sequences (release 2020-03) with known contaminants from MaxQuant and the cRAP repository. Trypsin was set as the proteolytic enzyme, allowing for up to two missed cleavages. Carbamidomethylation of cysteines was defined as a fixed modification, while oxidation of methionine and N-terminal acetylation were considered variable. The false discovery rate (FDR) was controlled below 1% at both peptide and protein levels. The “match between runs” feature was enabled to improve peptide identification across samples. The results files of Maxquant were analysed with Perseus version 1.6.14.0 (Tyanova et al., 2016). Contaminants and reverse proteins were eliminated. The data were transformed to Log 2 and only proteins with at least 3 valid values in at least one group were kept. Student T tests was applied to determine proteins of Interest (POI), with a pvalue <0.04 and Fc<-1.2 or Fc>1.2.

For the SETDB1-TGFβ secretome, MS raw files were processed using PEAKS Online X (build 1.8, Bioinformatics Solutions Inc.). Data were searched against the SwissProt database (https://www.expasy.org/resources/uniprotkb-swiss-prot). Parent mass tolerance was set to 20 ppm, with fragment mass tolerance at 0.05 Da. Specific tryptic cleavage was selected and a maximum of 2 missed cleavages was authorized. For identification, the following post-translational modifications were included: Acetylation (Protein N-term), oxidation, deamidation as variables and carbamidomethylation as fixed. Identifications were filtered based on a 1% false discovery rate threshold at both peptide and protein group levels. Label free quantification was performed using the PEAKS Online X quantification module, allowing a mass tolerance of 10 ppm, a CCS error tolerance of 0.02 and a 0.5-min retention time shift tolerance for match between runs. Protein abundance was inferred using the top N peptide method and TIC was used for normalization. Multivariate statistics on proteins were performed using Qlucore Omics Explorer 3.8 (Qlucore AB, Lund, *SWEDEN*). A positive threshold value of 1 was specified to enable a log2 transformation of abundance data for normalization *i.e.* all abundance data values below the threshold will be replaced by 1 before transformation. The transformed data were finally used for statistical analysis *i.e.* evaluation of differentially present proteins between two groups using a Student’s bilateral t-test. A p-value better than 0.01 was used to filter differential candidates. Proteomic data were analyzed using the functional annotation tool DAVID (Database for Annotation, Visualization, and Integrated Discovery, https://davidbioinformatics.nih.gov/home.jsp) (Sherman et al., 2022).

### Statistical analyses

Statistical analyses were carried out using Excel or Graphpad Prism (version 8.0, GraphPad Software Inc.). Data are represented as means ± SEM or mean ± SD as described in the figure legend. Two-tailed *t* test was used for statistical analysis; \**P* < 0.05, \*\**P* < 0.01, and \*\*\**P* < 0.001.

## DATA AVAILABILITY

All data needed to evaluate the conclusions in the paper are present in the paper and/or the Supplementary Materials. Data corresponding to all the experiments described in this study are deposited on ProteomeXchange Consortium via the PRIDE partner repository.

## ACKNOWLEDGEMENTS

We thank members of the Ait-Si-Ali lab and Epigenetics and Cell Fate department for helpful discussions during the group and department meetings and critical reading of the manuscript. We thank Dr Vincent Mouly and Kamel Mamchaoui from the Institute of Myology (Paris, France). We warmly thank Dr Paola Spessotto, the MyoLine platform for immortalization of human cells, the Myobank-AFM biobank from the Institute of Myology, and the biobank of the London MRC Neuromuscular Translational Research Centre and the Biomedical Research Centre of Great Ormond Street Hospital for Children in London for generous sharing of biological material. We thank Johanna Bruce and Philippe Chafey from Proteom’IC core facility of Institut Cochin in Paris for data acquisition and analyses of the proteomic data. The ProteoSeine platform of Institut Jacques Monod thanks Région Ile de France (SESAME 2021 EX061040), CNRS and Université Paris Cité for financial support.

## AUTHOR CONTRIBUTIONS

Conceptualization and experimental design: EN, MM, CT, SAS; Data acquisition and analyses: MZ, LM, AG, AB, GC, MB, JO, VL, VJ, MM; Figure composition and editing: MZ, LM, VJ, EN, MM, CT, SAS; Original draft: MZ, LM, VJ, EN, MM, CT and SAS; Writing – Review & Editing: MZ, LM, AB, VJ, EN, MM, CT, SAS; Supervision: EN, MM, CT and SAS; Funding Acquisition: AB, EN, CT, SAS. All authors have reviewed and approved the final manuscript for publication.

## SUPPLEMENTARY MATERIALS

Table S1: ANOVA analysis of all proteins identified from DMD secretome with SETDB1 KD and TGFβ treatment

Table S2: T-test analysis with siSETDB1 vs siSETDB1 +TGFβ (p-value <0.01)

Table S3: T-test analysis with siCTL vs siSETDB1 (p-value <0.01)

Table S4: T-test analysis with siCTL vs siCTL+TGFβ (p-value <0.01)

Table S5: T-test analysis with siCTL+TGFβ vs siSETDB1+TGFβ (p-value <0.01) Tables will be provided in the final version of the paper.

---

**Figure S1.**
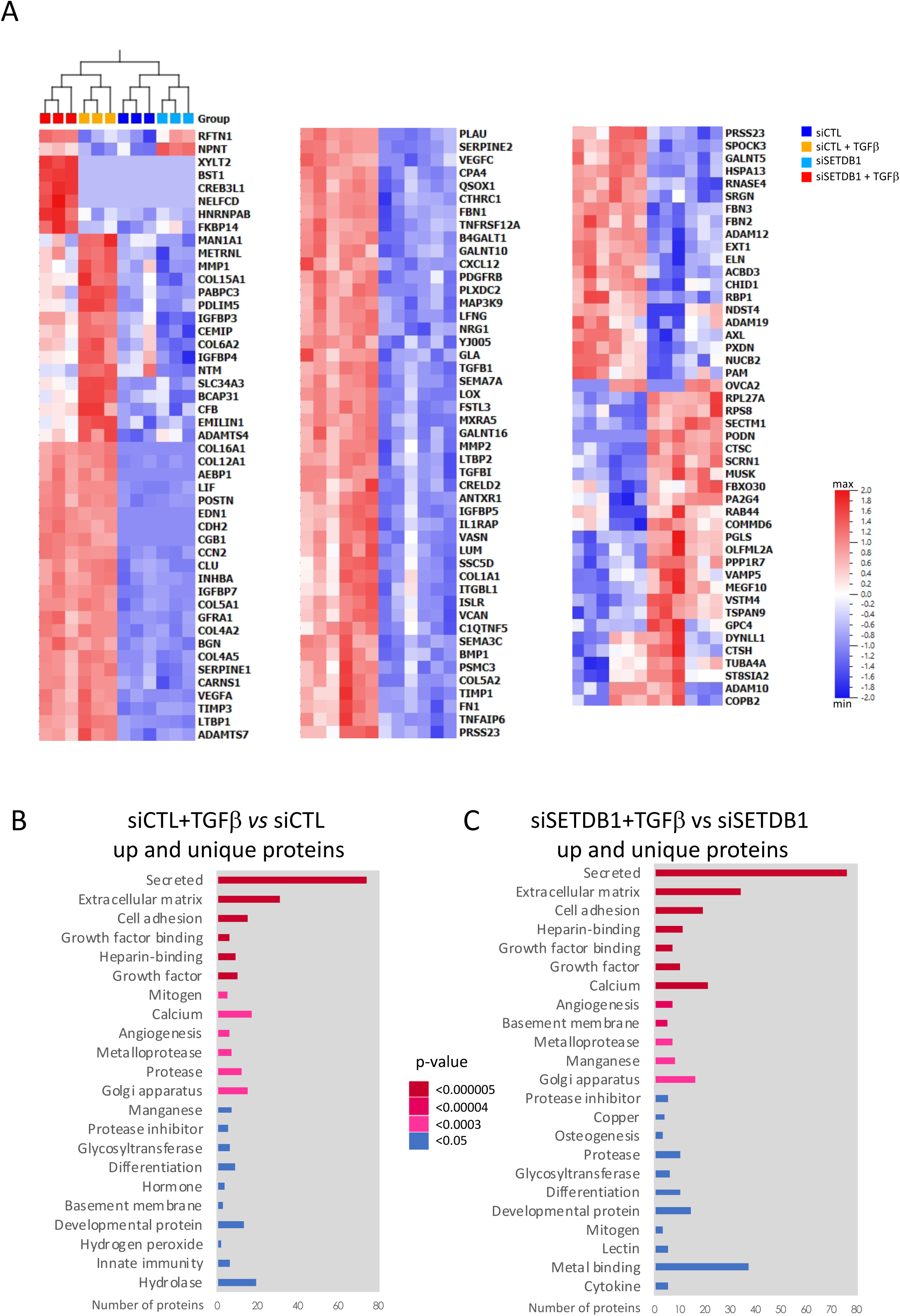
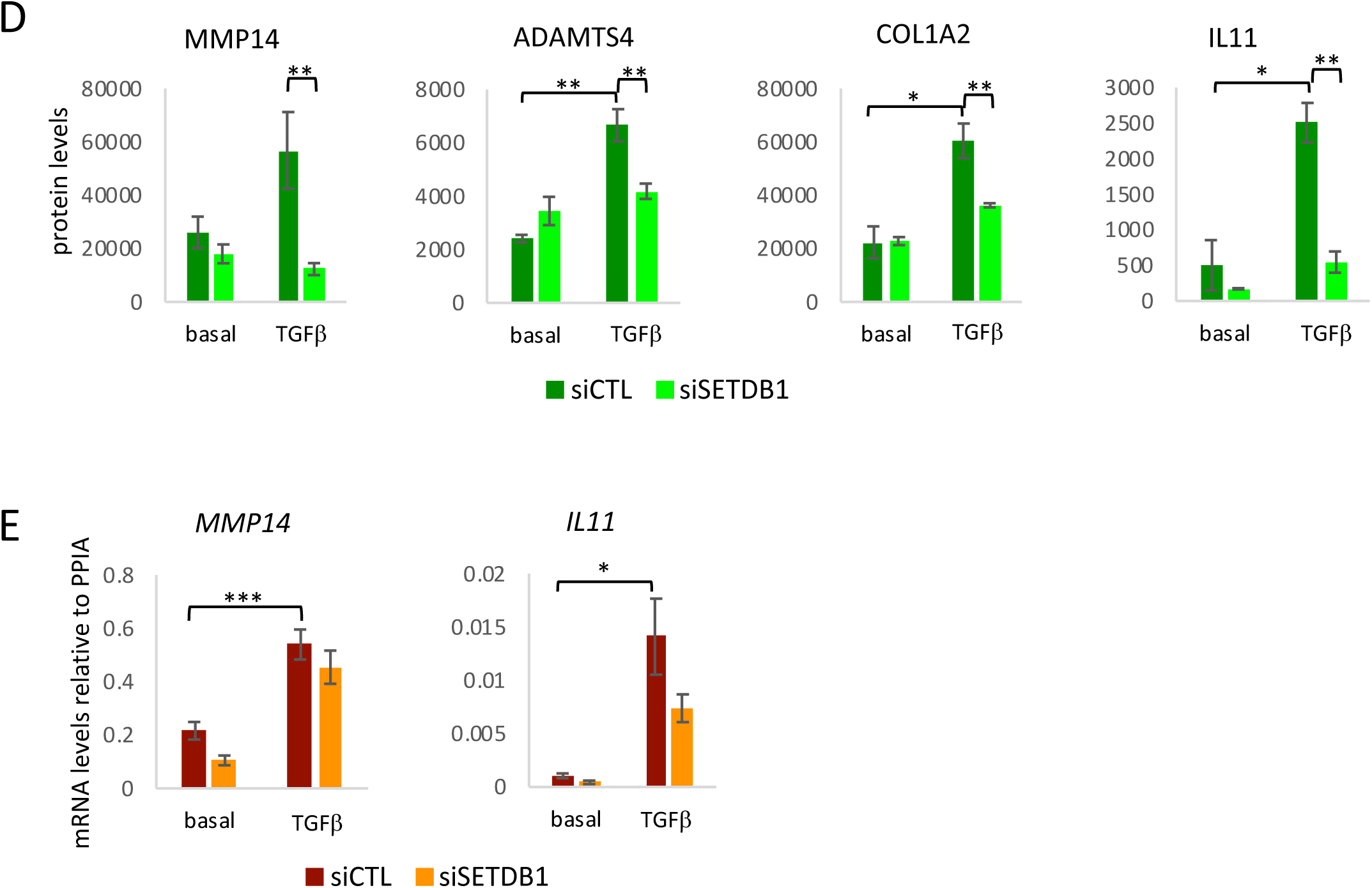
Detailed proteomic analysis of the TGFβ- and SETDB1-regulated secretome of DMD myotubes and analysis of SETDB1-dependent targets. **A.** Complete heatmap (ANOVA, p-value < 0.001) from the secretome analysis described in Fig.1. **B.** Davidbioinformatics functional annotation chart (Sherman et al., 2022) using uniprotKB keywords (biological process, cellular component, molecular function and ligand) using up and unique proteins from siCTL+TGFβ versus siCTL comparison. **C.** Davidbioinformatics functional annotation chart (Sherman et al., 2022) using uniprotKB keywords (biological process, cellular component, molecular function and ligand) using up and unique protein from siSETDB1+TGFβ versus siSETDB1 comparison. **D.** MS data analysis of MMP14 (matrix metalloproteinase-14), ADAMTS4 (a disintegrin and metalloproteinase with thrombospondin motif 4), COL1A2 (collagen type I alpha 2), IL11 (interleukin 11). **E.** RTqPCR of *MMP14* and *IL11* (N=3, DMD #1). For all panels data are represented as average -/+ SEM *p<0.05; **p<0.01; ***p<0.001 unpaired Student’s t test).

**Figure S2.**
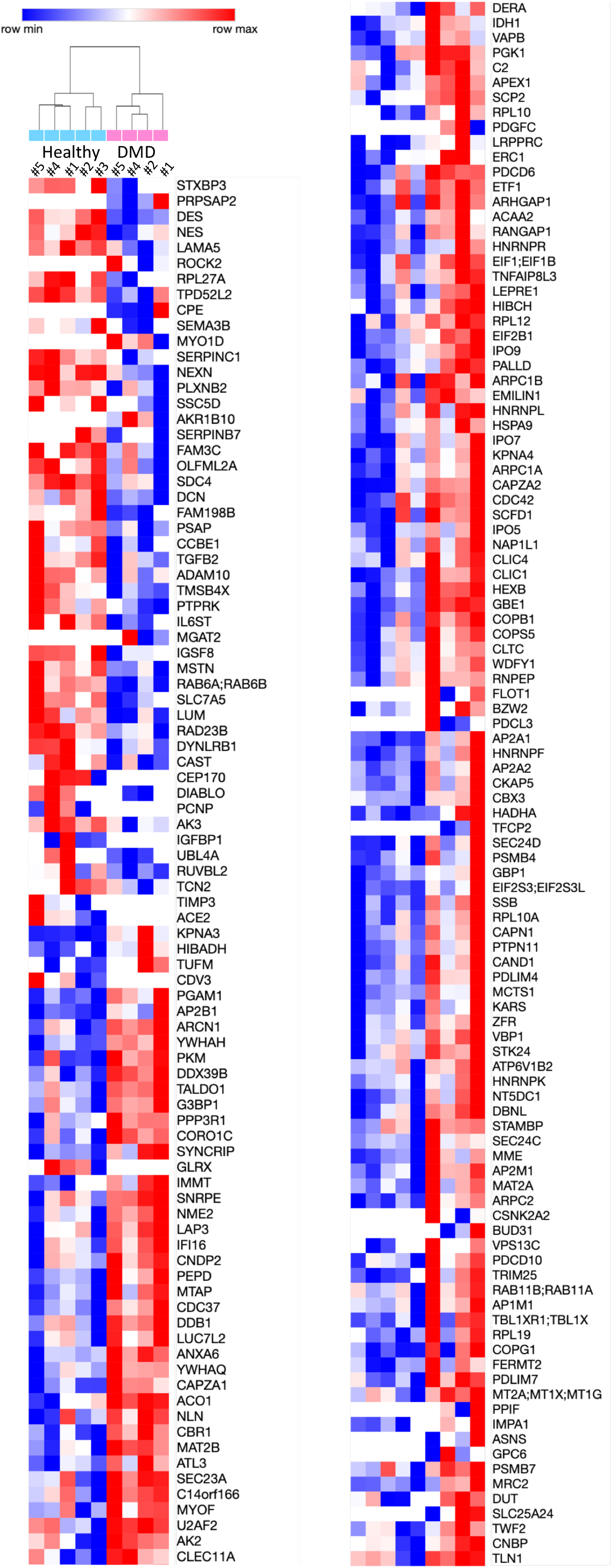
Detailed proteomic analysis of the DMD myotubes. Detailed heatmap from the secretome analysis described in Fig.2D.

**Figure S3.**
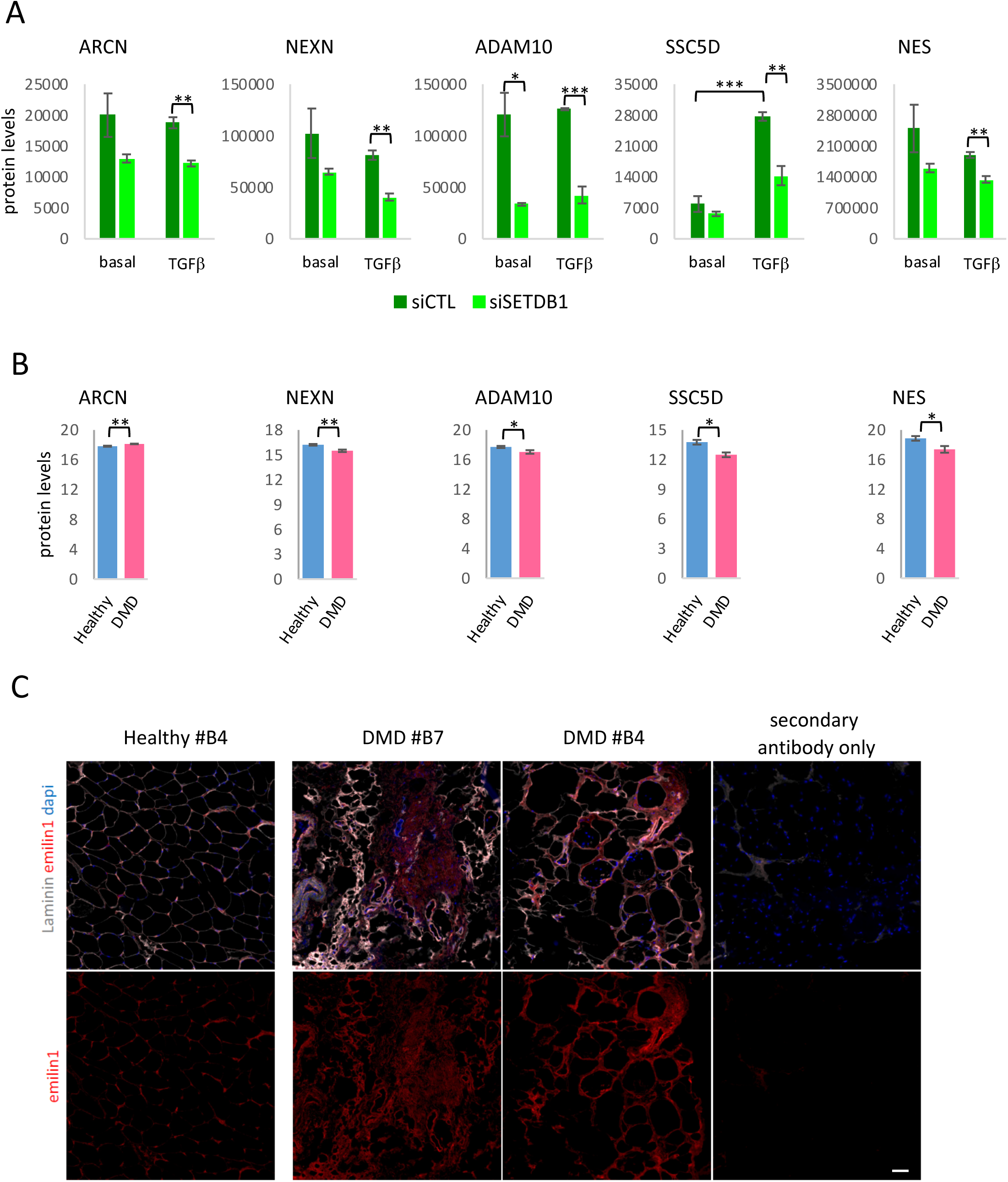
Common targets of the basal DMD and TGFβ–SETDB1 secretome. **A & B.** MS data analysis of ARCN (Coatomer subunit delta), NEXN (Nexilin), ADAM10 (Disintegrin and metalloproteinase domain-containing protein 10), SSC5D (Soluble scavenger receptor cysteine-rich domain-containing protein), NES (Nestin) with SETDB1 KD +/- TGFβ (**A**) and between healthy and DMD secretome (**B**). Data are represented as average +/-SEM *p<0.05; **p<0.01; ***p<0.001 unpaired Student’s t test. **C.** Immunostaining of EMILIN1 (Ab As556) (red), Laminin, and Dapi (blue) on human muscle biopsies of healthy and DMD patients, scale bar = 50µm.

**Figure S4.**
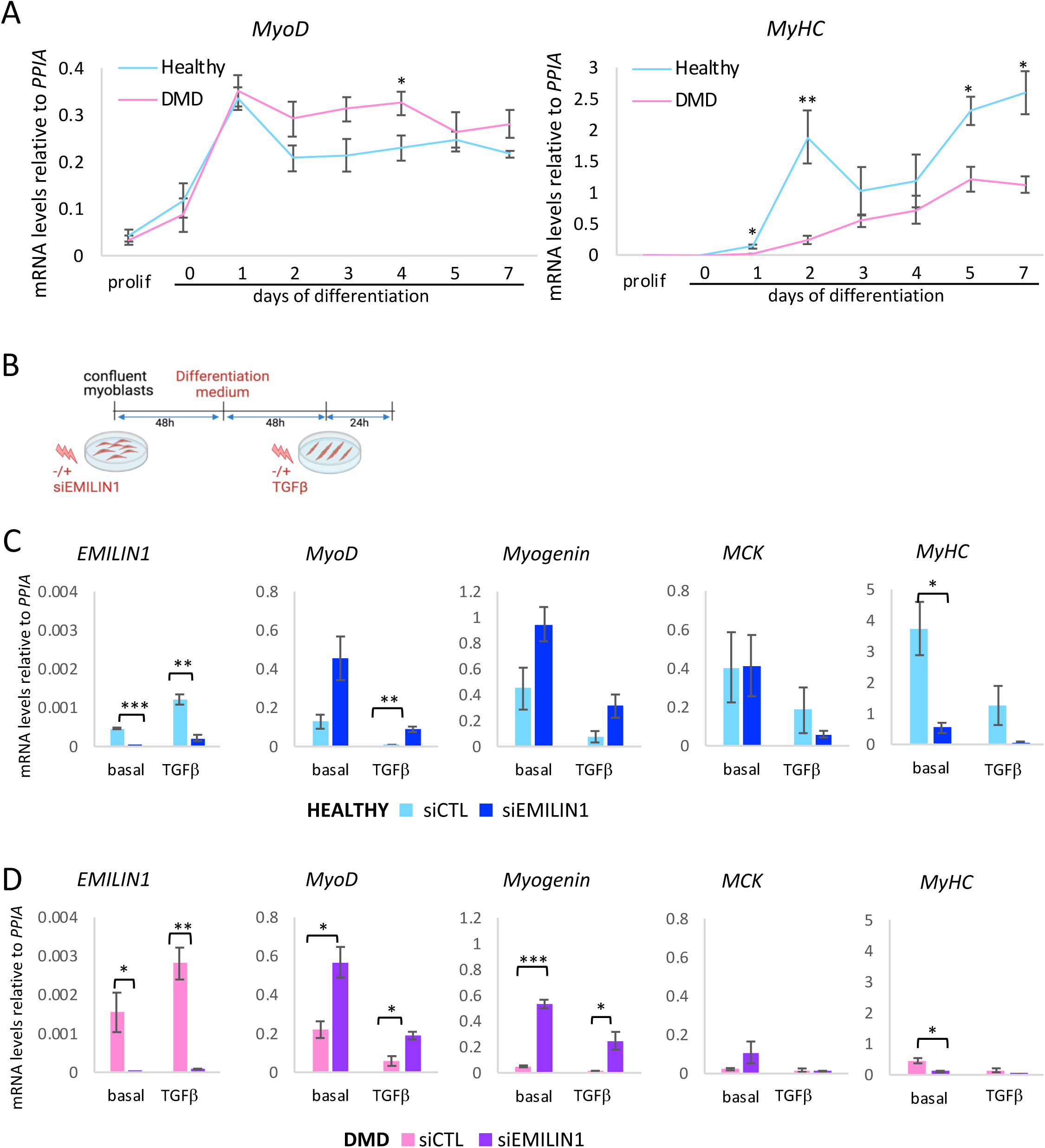
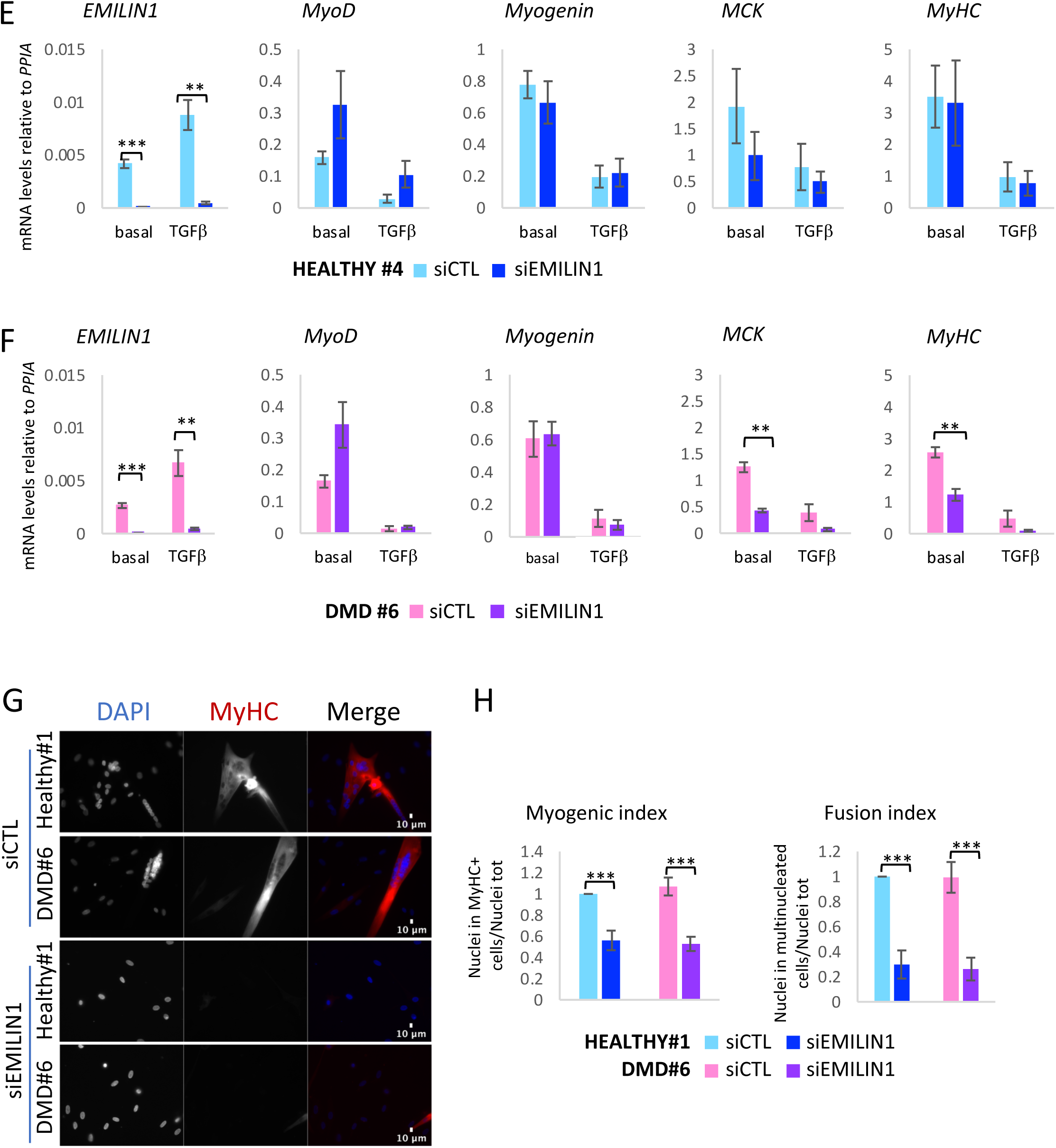
EMILIN1 KD regulates myogenic differentiation by promoting early markers and inhibiting late differentiation in healthy and DMD myoblasts and myotubes. **A.** RTqPCR of MyoD and MyHC in healthy and DMD proliferating myoblasts and differentiated myotubes (N=4, Healthy #1, DMD #1). **B.** Scheme of experimental design. Proliferating myoblasts were transfected with siRNAs scrambled (siCTL) or against EMILIN1 (siEMILIN1) for 2 days. Next, they were put in differentiation medium for 2 days and then treated or not with TGFβ at 20ng/mL for 24h. **C-D.** RTqPCR of EMILIN1 and of early (MyoD and Myogenin) and late (MCK and MyHC) myogenic markers in healthy (C) and DMD (D) myotubes with siCTL or siEMILIN1, +/- TGFβ (N=3, Healthy #1, DMD #1). **E-F.** RTqPCR of EMILIN1 and of early (MyoD and Myogenin) and late (MCK and MyHC) myogenic markers in healthy #4 (E) and DMD #6 (F) myotubes, collected accordingly to the experimental design Figure 3B (N=3). **G.** Immunofluorescence of healthy #1 and DMD #6 myotubes with siCTL or siEMILIN1 of MyHC (red) and nuclei staining with DAPI (blue). Scale bar 10μm. A representative field is shown for each condition. **H.** Quantification of myogenic and fusion index (N=7, Healthy #1, DMD #6). For all panels, data are represented as average -/+ SEM *p<0.05; **p<0.01; ***p<0.001 unpaired Student’s t test.

**Figure S5.**
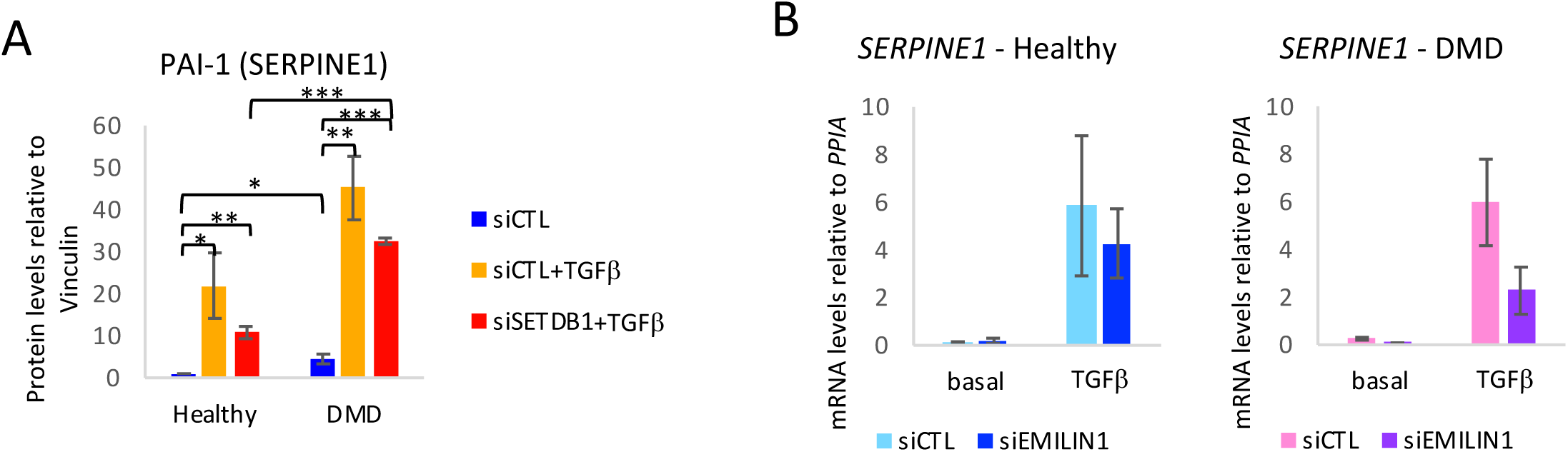
EMILIN1 KD reduces the expression of the TGFβ- and SETDB1- dependent fibrotic marker SERPINE1 in healthy and DMD myoblasts. **A.** Quantification of the western blot 5B normalized on Vinculin (N=4, DMD #1). For all panels, data are represented as average -/+ SEM *p<0.05; **p<0.01; ***p<0.001 unpaired Student’s t test. **B.** RTqPCR of SERPINE1 in healthy and DMD myoblasts +/- siEMILIN1, +/- TGFβ as in figure S4B (N=3, Healthy #1, DMD #1). For all panels, data are represented as average -/+ SEM *p<0.05; **p<0.01; ***p<0.001 unpaired Student’s t test.

